# Structural plasticity of axon initial segment in spinal cord neurons underlies inflammatory pain

**DOI:** 10.1101/2022.07.24.501281

**Authors:** Yaki Caspi, Michael Mazar, Yishai Kushnir, Yoav Mazor, Ben Katz, Shaya Lev, Alexander M Binshtok

## Abstract

Activity-dependent structural plasticity of axon initial segment (AIS) regulates neuronal excitability, thus fine-tuning neuronal and overall network output. Here using behavioral, immunohistochemical, electrophysiological and computational approaches, we describe the structural plasticity of AIS in rat’s superficial spinal cord dorsal horn (SDH) neurons, which underlies inflammatory pain. We show an inflammation-mediated distal shift of the AIS away from the soma in inhibitory but not excitatory SDH neurons, concomitant with the peak of inflammatory pain. This AIS translocation was accompanied by a decrease in excitability of the inhibitory neurons. Following recovery from inflammatory hyperalgesia, the AIS location and neuronal excitability reversed to baseline levels. The computational model of SDH inhibitory neurons predicts that the distal shift of AIS is sufficient to decrease the intrinsic excitability of these neurons. Our results provide evidence of differential inflammation-mediated AIS plasticity, reducing the excitability of inhibitory but not excitatory SDH neurons and contributing to inflammatory hyperalgesia.

## Introduction

Tissue inflammation or injury induces an increase in the excitability of primary afferent neurons (Binshtok et al., 2008; Djouhri et al., 2006; Goldstein et al., 2019; Gudes et al., 2015; Jin and Gereau, 2006; Sorkin et al., 1997) amplifying their input towards the nociceptive CNS networks and resulting in enhanced pain sensitivity, namely, hyperalgesia (Barkai et al., 2020; Goldstein et al., 2019; Yarmolinsky et al., 2016). The “first stations” processing the inputs from primary afferents are the intricate neuronal networks of superficial laminae of the dorsal spinal cord (SDH) (Basbaum et al., 2009). These pain-related neuronal networks are composed of local excitatory and inhibitory neurons converging into projection neurons, which relay the information to higher brain centers, thus forming the sensation and perception of pain (Braz et al., 2014; Todd, 2010; Wercberger and Basbaum, 2019). It is well documented that inflammation and injury-mediated changes in the primary afferent input lead to an increase in excitatory synaptic transmission towards local interneurons and projection neurons (Baba et al., 1999; Kohno et al., 2003; Li et al., 2015; Shevchuk et al., 2017). In other neuronal systems, enhanced synaptic bombardment and increased neuronal activity trigger an adaptive decrease in intrinsic neuronal excitability, hence regulating neuronal output (Grubb and Burrone, 2010; Jamann et al., 2021; Lezmy et al., 2020, 2017). The adaptive changes tuning the activity of pain-related SDH neurons, thus controlling nociception and pain, have not been described before.

It has been demonstrated that the plasticity of axon initial segment (AIS) is an essential mechanism for tuning neuronal and hence network output in response to changes in input (Buffington and Rasband, 2011; Grubb et al., 2011; Huang and Rasband, 2018; Kole and Brette, 2018; Kole and Stuart, 2012). The AIS, is located at the proximal axon and harbors a dense array of voltage-gated channels and a unique repertoire of submembranous cytoskeletal scaffolding proteins, such as ankyrin G, clustering these channels together (Grubb et al., 2011; Ha and Rasband, 2019; Huang and Rasband, 2018). The high density of voltage-gated channels allows AIS to convert graded synaptic inputs into a zero-one binary output of action potential (AP) firing. AIS (Battefeld et al., 2014; Kole et al., 2008; Kole and Stuart, 2012). The variations in the voltage-gated channel composition at the AIS determine the pattern of neuronal firing, hence defining neuronal output (Bender and Trussell, 2012; Kole and Brette, 2018; Kole and Stuart, 2012; Lorincz and Nusser, 2008; Sasaki, 2013). Additionally, the structural properties of AIS, such as the AIS size and location relative to soma, determine the activity pattern of neurons (Huang and Rasband, 2018; Kuba et al., 2006; Lorincz and Nusser, 2008). Notably, the size and position of individual AIS are dynamic and modified in response to physiological and pathological changes in neuronal activity (Galliano et al., 2021; Grubb and Burrone, 2010; Jamann et al., 2021, 2018; Kuba et al., 2010; Leterrier, 2018; Lezmy et al., 2020). This type of structural AIS plasticity affects neuronal excitability, thus fine-tuning neuronal output and the overall neuronal network response to changes in input (Grubb and Burrone, 2010; Jamann et al., 2021; Kuba et al., 2010; Lezmy et al., 2017; Rotterman et al., 2021). Sensory deprivation, for example, leads to AIS elongation and a proximal shift of its position towards the soma, resulting in an increase in neuronal excitability (Jamann et al., 2021; Kuba et al., 2010). On the other hand, sensory enrichment or pharmacologically-induced increase in neuronal activity produces a distal shift of AIS location and AIS shortening, leading to a decrease in neuronal excitability (Grubb and Burrone, 2010; Jamann et al., 2021; Lezmy et al., 2020, 2017). However, it is unknown whether inflammation or tissue injury-induced increase in input to SDH neurons induces plasticity of AIS, thus affecting the excitability of SDH neurons.

Here we studied the changes in the location of AIS in inhibitory and excitatory SDH neurons following hyperalgesic inflammation and detailed how these changes alter the intrinsic neuronal excitability. We showed that in normal conditions, the AIS in Pax2^+^ inhibitory SDH neurons is located closer to the soma than in Pax2^−^ excitatory neurons. We also showed that 24 h following injection of proinflammatory Complete Freund Adjuvant (CFA), when inflammatory hyperalgesia is at its peak, AIS in inhibitory but not in excitatory neurons shifts distally away from the soma. Using the patch-in-slice approach, we confirmed the inflammation-mediated change in AIS location in tonically-firing inhibitory neurons and demonstrated a significant increase in threshold current (rheobase) of inhibitory neurons. The excitability of excitatory neurons was not affected. Furthermore, the changes in AIS location and the increase in threshold current of inhibitory neurons followed a timeline concomitant with the development of hyperalgesia, such that the AIS location and threshold current returned to baseline values as the animals recovered from hyperalgesia. To better correlate the distal shift in AIS locations to the changes in the excitability of inhibitory neurons, we built a multicompartmental model of SDH inhibitory neurons, which predicted that distal shift in AIS is sufficient to produce an increase in threshold current. Our results suggest that inflammation triggers plasticity of the AIS selectively in inhibitory SDH neurons. The inflammation-mediated distal shift of AIS and the resulting decrease in inhibitory neurons’ excitability could contribute to an increase in output from the pain-related SDH network, thus facilitating inflammatory pain.

## Results

### AIS in inhibitory but not excitatory SDH neurons shifts away from the soma during inflammatory hyperalgesia

Inflammation or tissue injury produces an increase in the gain of primary nociceptive neurons (Barkai et al., 2019; Binshtok et al., 2008; Goldstein et al., 2019; Hucho and Levine, 2007; Lennertz et al., 2012), thus enhancing their output towards the second-order spinal cord neurons (Baba et al., 1999; Shevchuk et al., 2017). It has been demonstrated in various neuronal systems that abrupt changes in either intrinsic or synaptic activity lead to changes in size and location of AIS, thus inducing adaptive changes in neuronal excitability (Grubb and Burrone, 2010; Jamann et al., 2021; Kuba et al., 2010; Lezmy et al., 2020, 2017). Here we studied the effect of inflammation on the intrinsic excitability and AIS plasticity in excitatory and inhibitory spinal cord neurons at the superficial dorsal horn (SDH). First, we examined whether inflammation leads to AIS plasticity using immunohistochemical staining of spinal cord slices from rats 24 - 48 h after injecting CFA to the left paw. We chose this time window since our behavioral experiments demonstrated that CFA-induced inflammatory hyperalgesia peaks at this time (Figure 1A, *red circles*). The AIS were detected using staining for ankyrin-G, a protein composing the AIS (Kordeli et al., 1995), and the inhibitory interneurons were identified using staining for Pax2, a transcription factor promoting inhibitory neurons cell fate (Punnakkal et al., 2014). Due to the complex axonal geometry of SDH neurons, we could not accurately measure the size of AIS in the spinal cord slices (*see Methods*) and therefore focused on studying the changes in AIS’ distance from the soma. We measured the AIS distance from the soma in SDH Pax2^+^ inhibitory and Pax2^−^ excitatory neurons (Punnakkal et al., 2014) on the ipsilateral side of the CFA injection (inflammatory conditions) and compared it to the distance of AIS from the soma on the contralateral side to CFA injection, in which no behavioral changes were observed (Figure 1A, *black squares*, control conditions).

**Figure 1.**
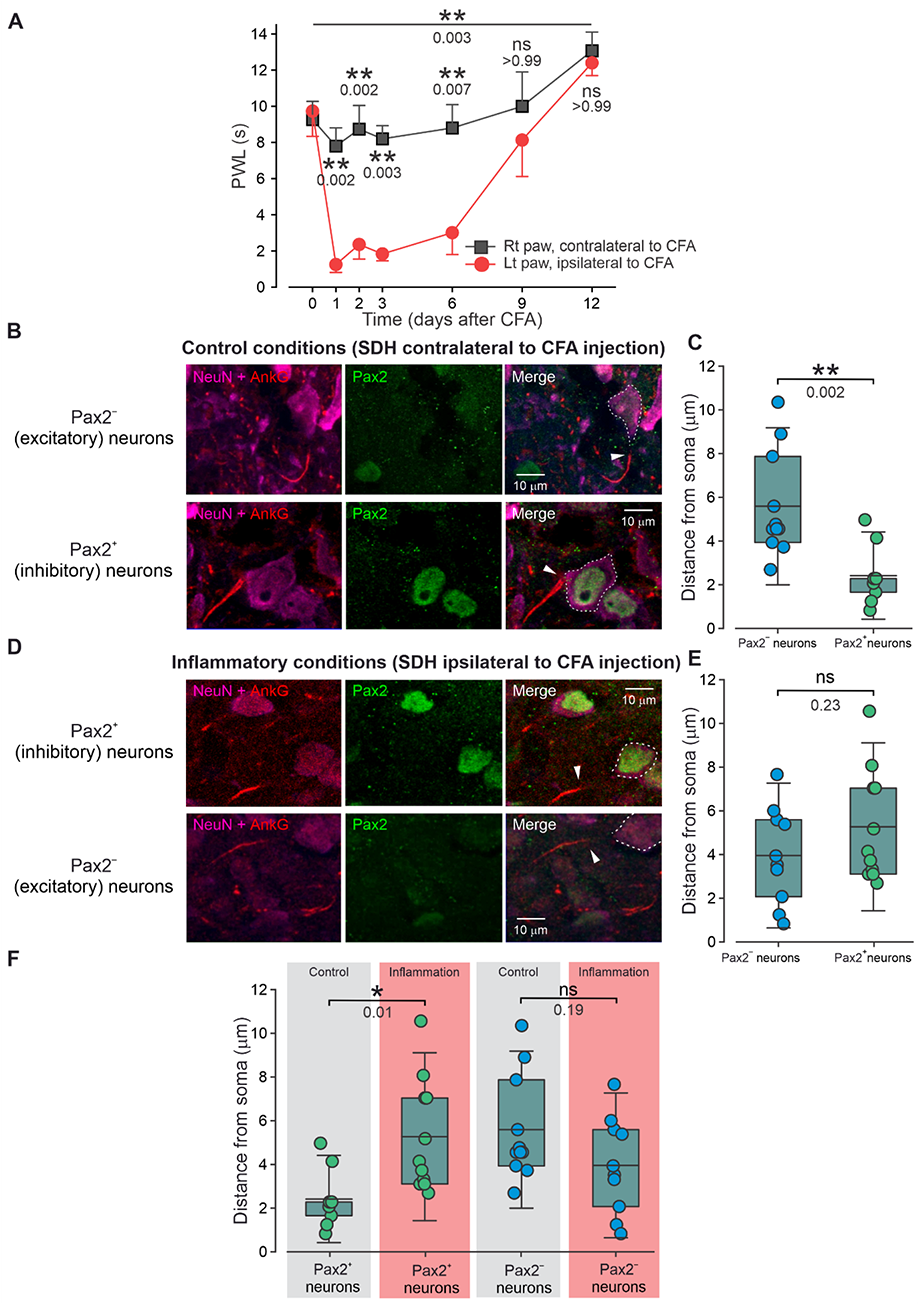
AIS in inhibitory but not excitatory SDH neurons shifts distally from the soma in inflammatory conditions. **A.** Changes in paw withdrawal latency (PWL) following CFA injection into the left paw measured in the left (ipsilateral) and the right (contralateral) paw. Timepoint “0” depicts the baseline measurements before CFA injection. n = 7 rats, ordinary two-way ANOVA with post hoc Bonferroni. **B.** Representative confocal images of SDH neurons stained for NeuN (*magenta*), AnkG (*red*) and Pax2 (*green*) and their merge (*right*) in control conditions. *The upper panels* show Pax2^−^ cell, and the *lower panels* show Pax2^+^ cells. The dotted lines on the right panels outline the soma border and are based on the NeuN staining. The arrow indicates the beginning of AIS based on the AnkG staining. **C.** Box plots and individual values of the distances of AIS from the soma measured in Pax2^−^ and Pax2^+^ neurons in control conditions. Each data point represents individual neurons. One neuron per rat was used: n _Pax2_^−^, _Naive_ = 11 neurons in 11 rats; n _Pax2_^+^, _Naïve_ = 9 neurons in 9 rats, Student *t*-test. **D.** Same as in *B* but showing representative Pax2^−^ and Pax2^+^ neurons in inflammatory conditions. **E.** Box plots and individual values of the distances of AIS from the soma measured in Pax2^−^ and Pax2^+^ neurons in inflammatory conditions. n _Pax2_^−^, _Inflammation_ = 10 neurons in 10 rats; n _Pax2_^+^, _Inflammation_ = 11 neurons in 11 rats, Student *t*-test. **F.** Data shown in *C* and *E* rearranged to compare the distances of AIS from the soma in Pax2^+^ and Pax2^−^ neurons between control (*contra, grey*) and inflammatory (*ipsi, light red*) conditions. n _Pax2_^+^, _Naïve_ = 9 neurons in 9 rats; n _Pax2_^+^, _Inflammation_ = 11 neurons in 11 rats; n _Pax2_^−^, _Naïve_ - 11 neurons in 11 rats; n n _Pax2_^−^, _Inflammation_ - 10 neurons in 10 rats; Ordinary one-way ANOVA with posthoc Bonferroni.

Surprisingly, we found that in control conditions, AIS in inhibitory Pax2^+^ neurons were located ∼2 μm (2.4 ± 1.3 μm, n=9 neurons in 9 rats) from the soma, which was significantly closer to the soma than in excitatory Pax2^−^ neurons (5.6 ± 2.4 μm, n=11 neurons in 11 rats, p = 0.002, Student *t*-test, Figure 1B, C).

Notably, in inflammatory conditions, the AIS in the Pax2^+^ inhibitory neurons assumed a more distal location of ∼5 μm (5.3 ± 2.6 μm, n=11 neurons in 11 rats) from the soma (Figure 1D, E). The location of AIS in inhibitory neurons in the inflammatory conditions was significantly more distal than the location of inhibitory neurons’ AIS under control conditions (Figure 1F). This shift in AIS was selective for inhibitory neurons since the AIS distance from the soma was similar in Pax2^−^ excitatory neurons in inflammatory and control conditions (Figure 1F). This data suggests that AIS in inhibitory neurons shifts distally from soma during inflammatory hyperalgesia.

### Tonically-firing SDH inhibitory neurons exhibit mostly monophasic phase plots in naïve conditions and largely bi-phasic phase plots in inflammatory conditions

Since AIS location defines the threshold current of neurons (Grubb and Burrone, 2010; Rotterman et al., 2021), the shift of AIS in inhibitory neurons suggests that inhibitory neurons might change their excitability under inflammatory conditions. We have measured the excitable properties of SDH inhibitory and excitatory neurons using patch-in-slice recordings from spinal cord slices obtained from naïve rats and rats 24 hours after CFA injection into the left paw (inflammatory conditions). We distinguished between inhibitory and excitatory neurons by characterizing their firing pattern, using stimulation with 500 ms increasing current pulses (Figure 2A, Supplementary Figure 1). It has been shown that the absolute majority of the tonically-firing SDH neurons (neurons that fire multiple APs at threshold depolarization, Figure 2A, Supplementary Figure 1A, *right*) are inhibitory (Punnakkal et al., 2014). Most of the delayed-firing SDH neurons (neurons in which the first AP occurs with a delay of about hundreds of ms, Figure 2A, Supplementary Figure 1A, *left*) are excitatory (Punnakkal et al., 2014). To assure that neurons do not change their firing patterns with increased stimulation current, we characterized the firing properties of the SDH neurons along different amounts of current injections (Figure 2A, Supplementary Figure 1). None of the recorded delayed-firing (n=11 neurons, 11 slices, 6 rats) or tonically-firing neurons (n=13 neurons, 13 slices, 7 rats) changed their firing or delay to the first AP with increasing stimulation (Supplementary Figure 1B, C). Consequently, we considered the neurons exhibiting tonic firing patterns as inhibitory and neurons with delayed firing patterns - excitatory (Figure 2A).

**Figure 2.**
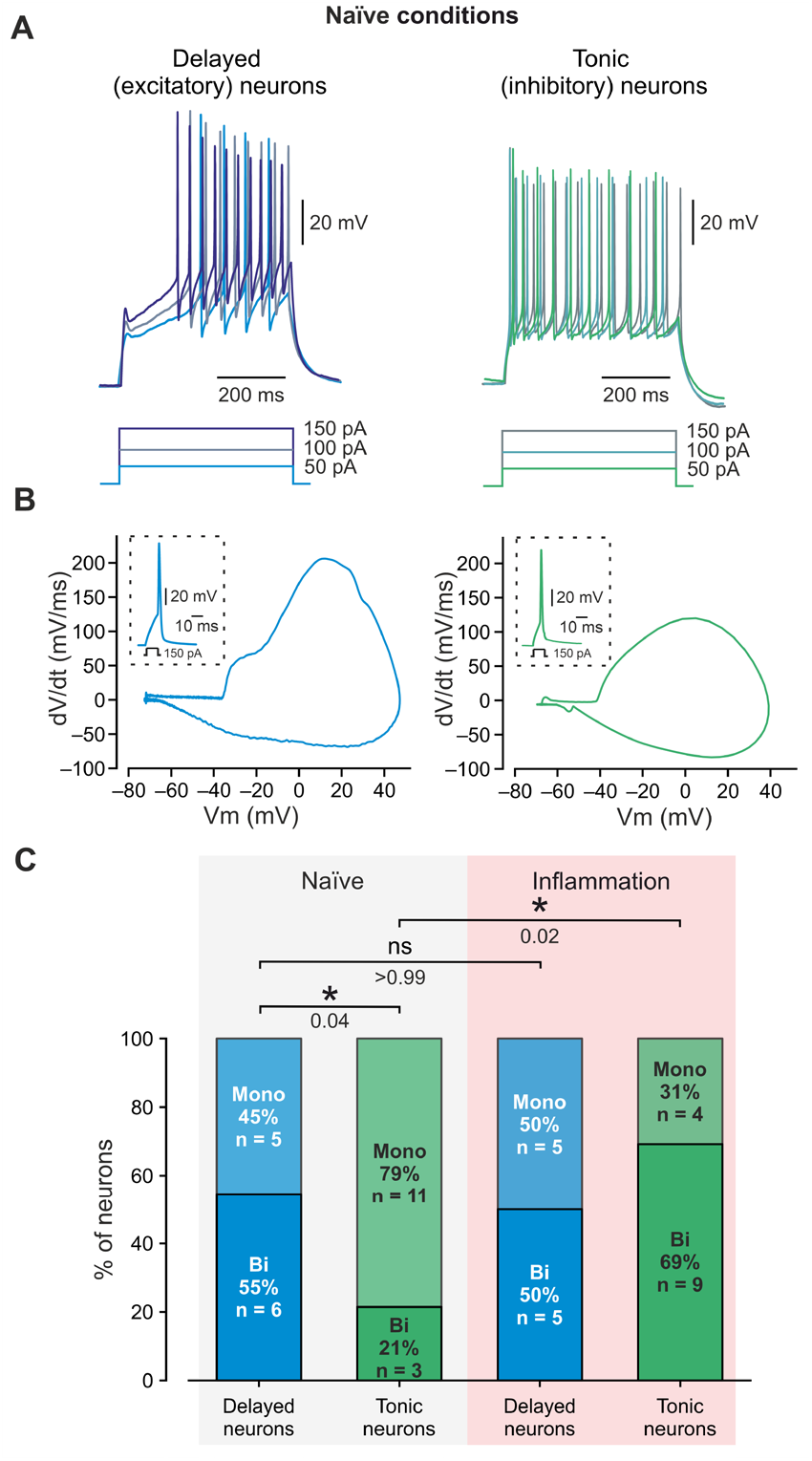
In inflammatory conditions, a majority of tonically firing SDH neurons switch from a monophasic to a bi-phasic phase plot. **A.** Representative traces of current-clamp recordings from two SDH neurons showing delayed-firing (“delayed,” *left, shades of blue*) and tonically-firing (“tonic,” *right, shades of green*) patterns recorded following increasing current steps of 50, 100 and 150 pA. Note that the delayed-and tonically-firing patterns remained stable along with increasing stimulation currents (see also Supplementary Figure 1). Representative of 11 delayed neurons (from 11 slices, 7 rats) and 13 tonic neurons (13 slices, 7 rats). **B.** Examples of bi-phasic (*left, blue*) and monophasic (*right, green*) phase plots of the rate of change in the membrane potential (d*V*/d*t*) during an AP (shown in *insets*) vs. membrane potential recorded from delayed-firing (*left*) and tonically-firing (*right*) SDH neurons, respectively. **C.** 100% stacked column graphs depicting the percentage of monophasic and biphasic phase plots recorded from delayed (*blue*) and tonically-firing (*green*) neurons in naïve and inflammatory conditions. Note the increase in the percentage of the biphasic phase plots in tonically-firing neurons in inflammatory conditions and no change in the percentage of biphasic phase plots in delayed-firing neurons. n _Delayed, Naïve_ = 11 neurons, 11 slices, 6 rats; n _Tonic, Naïve_ = 14 neurons, 14 slices, 7 rats; n _Delayed, Inflammation_ = 10 neurons, 10 slices, 5 rats; n _Tonic, Inflammation_ = 13 neurons, 13 slices, 6 rats, Fisher’s exact test.

Moreover, it has been demonstrated that the relative distance of AIS from the soma could be predicted by the shape of the phase plot of the AP (Goethals and Brette, 2020; Kole and Brette, 2018; Lezmy et al., 2017; Telenczuk et al., 2017). AP initiated at the AIS located far away from the soma gives rise to a stereotypical “humped” bi-phasic phase plot, with a distinguishable “early” phase, reflecting AP generation at the AIS, while AP generated at the AIS located close to the soma produces a monophasic phase plot since voltage changes at the AIS are masked by voltage changes at the somatic membrane (Goethals and Brette, 2020; Grubb and Burrone, 2010; Kole and Brette, 2018; Lezmy et al., 2017). Therefore, we performed electrophysiological recordings of SDH neurons from spinal cord slices of naïve rats, and after characterizing the neuronal firing patterns (Figure 2A, Supplementary Figure 1), we computed the phase plots of the single action potential of tonically-firing and delayed-firing SDH neurons (Figure 2B). We found that tonically-firing neurons exhibit a significantly higher number of monophasic phase plots than the delayed-firing neurons (11 out of 14 tonically firing neurons vs. 5 out of 11 delayed firing neurons; p = 0.04, Fisher’s exact test; Figure 2C, *Naïve*). Next, we examined SDH neurons from spinal cord slices under inflammatory conditions, i.e., at the ipsilateral side of CFA-treated rats. We found that under inflammatory conditions, tonically-firing SDH neurons possessed a significantly lower number of monophasic and a significantly higher number of bi-phasic phase plots (4 out of 13 neurons, p = 0.02, Fisher’s exact test; Figure 2C) when compared to naïve animals. No change in the proportion of mono- and bi-phasic phase plots was observed in delayed-firing SDH neurons (Figure 2C). These results suggest that the AIS of tonically firing inhibitory SDH neurons predominantly assume an adjacent-to-soma localization in naïve animals, while inflammation promotes a shift of the AIS to assume a more distant location. Moreover, the AIS location in delayed-firing excitatory SDH neurons is unaffected by inflammation.

Collectively, our results suggest that (*1*) in normal conditions, the location of AIS in inhibitory neurons is closer to the soma than in excitatory neurons and (*2*) AIS in inhibitory neurons is shifted distally from the soma following hyperalgesic inflammation. A distal AIS shift in inhibitory neurons implies that these neurons should demonstrate a change in their intrinsic excitable properties following inflammation.

### The threshold current of tonically-firing SDH neurons increases in hyperalgesic conditions

Theoretical (Goethals and Brette, 2020) and experimental data (Grubb and Burrone, 2010; Rotterman et al., 2021) suggest that a distal shift of the AIS from the soma affects the intrinsic excitability properties of neurons. Accordingly, we examined the intrinsic excitability properties of tonically- and delayed-firing SDH neurons under naïve and inflammatory conditions (Figure 3 **and** Table 1). We first examined the threshold current (rheobase, *see Methods*), the minimal current required for AP generation, since a large body of work has associated a shift in AIS location with changes in threshold current (Grubb and Burrone, 2010; Rotterman et al., 2021). We found that the threshold current of tonically firing SDH neurons in inflammatory conditions is significantly higher than in tonically firing neurons recorded from naïve rats (Figure 3A, B).

**Figure 3.**
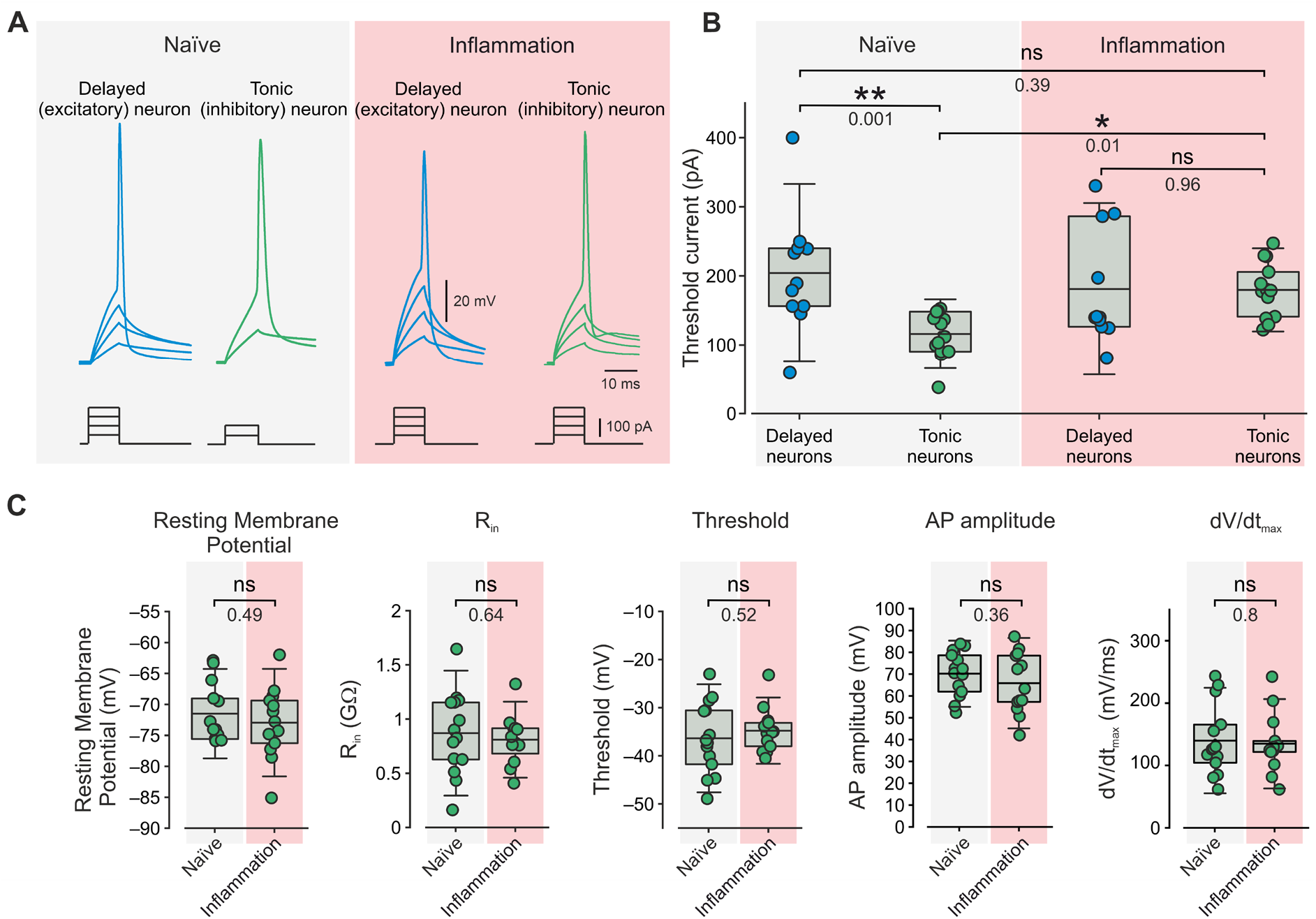
Inflammation leads to an increase in threshold current selectively in tonically firing SDH neurons. **A.** Representative traces of membrane voltage responses to 10 ms current steps recorded from delayed-firing (“delayed,” *blue*) and tonically-firing (“tonic,” *green*) neurons in naïve and inflammatory conditions. Note that AP in tonically-firing neurons is evoked by a substantially smaller current in naïve conditions than in inflammatory conditions. Representative of 11 delayed neurons (from 11 slices, 7 rats) and 13 tonic neurons (13 slices, 7 rats). **B.** Box plots and individual values of the threshold current assessed from delayed- (*blue*) and tonically-firing (*green*) neurons in naïve and inflammatory conditions. n _Delayed, Naïve_ = 11 neurons, 11 slices, 6 rats; n _Tonic, Naïve_ = 13 neurons, 13 slices, 7 rats; n _Delayed, Inflammation_ = 11 neurons, 11 slices, 5 rats; n _Tonic, Inflammation_ = 14 neurons, 14 slices, 6 rats, Ordinary one-way ANOVA with posthoc Bonferroni. **C.** Box plots and individual values of the excitability properties of tonically firing neurons in naïve and inflammatory conditions. n _Tonic, Naïve_ = 13 neurons, 13 slices, 7 rats; n _Tonic, Inflammation_ = 14 neurons, 14 slices, 6 rats, Student *t*-test.

**Table 1.** Comparison of excitability properties of tonically-firing and delayed-firing SDH neurons in naïve conditions, inflammatory conditions (24 - 48 h after injection of CFA) and recovery (9 - 14 days after injection of CFA).

| ***Tonically-firing neurons*** |  |  |  |  |  |  |
| --- | --- | --- | --- | --- | --- | --- |
|  | *Rheobase*<br>*(pA)* | *RMP (mV)* | *R<sub>in</sub> (GΩ)* | *Threshold*<br>*(mV)* | *AP*<br>*amplitude*<br>*(mV)* | *dV/dt<sub>max</sub>*<br>*(mV/ms)* |
| **Naïve**<br>(n = 13<br>neurons, 10<br>slices, 7 rats) | 116 ± 33 | -69.7 ± 8 | 0.87 ± 0.38 | -36.3 ± 7.5 | 70.2 ± 10 | 140 ± 56 |
| **Inflammation**<br>(n = 14<br>neurons, 11<br>slices, 6 rats) | 180 ± 40<br>***, p=0.0002<br>**, p=0.0013 | -72.9 ± 5.8<br>ns, p=0.69<br>ns, p>0.99 | 0.94 ± 0.52<br>ns, p>0.99<br>ns, p>0.99 | -34.8 ± 4.5<br>ns, p>0.99<br>ns, p=0.4 | 65.9 ± 14<br>ns, p=0.35 | 135 ± 48<br>ns, p=0.8 |
| **Recovery**<br>(n = 4 neurons,<br>4 slices, 3 rats) | 98 ± 25<br>ns, p>0.99 | -71.5 ± 4<br>ns, p>0.99 | 0.77 ± 0.33<br>ns, p>0.99 | -40.1 ± 4.2<br>ns, p=0.83 | 65.1 ± 14<br>ns, p>0.99<br>ns, p>0.99 | 122 ± 29<br>ns, p>0.99<br>ns, p>0.99 |
| ***Delayed-firing neurons*** |  |  |  |  |  |  |
| **Naïve**<br>(n = 11<br>neurons, 8<br>slices, 6 rats) | 204 ± 85 | -70.9 ± 4 | 0.83 ± 0.23 | -37.2 ± 13.7 | 62.4 ± 17 | 109 ± 44 |
| **Inflammation**<br>(n = 11<br>neurons, 9<br>slices, 4 rats) | 181 ± 83<br>ns, p=0.53 | -70.5 ± 2.5<br>ns, p=0.79 | 1.12 ± 0.64<br>ns, p=0.16 | -32.4 ± 8.7<br>ns, p=0.36 | 64.3 ± 13<br>ns, p=0.79 | 153 ± 62<br>ns, p=0.07 |

In addition to the distal shift in AIS location, an increase in threshold current could result from changes in resting membrane potential (RMP), decrease in input resistance (R_in_), or decreased availability of voltage-gated sodium channels, leading to a decrease in AP threshold (Bean, 2007) We, therefore, examined whether other excitable properties of SDH inhibitory neurons were changed following inflammation. No significant difference was found in RMP, R_in_, AP threshold, AP amplitude, and dV/dt_max_ (which reflects the availability of voltage-gated sodium channels (Bean, 2007; Jenerick, 1963)) between tonically-firing SDH neurons recorded from slices of naïve or CFA-treated rats (Figure 3C). These results suggest that the inflammation-induced increase in threshold current described here could be attributed at least in part to the inflammation-mediated shift in the AIS location.

Furthermore, we show that the inflammation-induced change in threshold current was selective to the tonically-firing neurons since no change in threshold current was detected in delayed firing neurons recorded in slices obtained from naïve or CFA-treated animals (Figure 3B). No changes in other excitability parameters in delayed firing neurons were observed as well (Table 1). Notably, we show that in naïve conditions, the threshold current of tonically-firing inhibitory neurons was significantly lower than that of delayed-firing excitatory neurons (Figure 3A, B). These results are in correlation with the difference in the AIS location between these neurons (Figure 1B, C). Moreover, in inflammatory conditions, when the AIS distance from the soma is similar in inhibitory and excitatory neurons (Figure 1D, E), the difference in threshold current between tonically- and delayed-firing neurons is annulled (Figure 2A, B), suggesting that AIS distance from the soma could determine the threshold current of a neuron.

### Inflammation-induced distal AIS shift and increase in threshold current reverse after recovery from hyperalgesia

To further establish a link between the shift of AIS location, changes in threshold current, and inflammatory hyperalgesia, we performed immunohistochemical staining and electrophysiological measurements of SDH neurons 9 - 12 days after injection of CFA, a time window at which CFA-induced hyperalgesia has subsided (Figure 1A). We performed immunohistochemical analysis of AIS location in Pax2^+^ neurons in the inflammatory conditions and compared it to AIS location in the control conditions. We show that at a time point when hyperalgesia subsides, the AIS of SDH Pax2^+^ inhibitory neurons is located significantly closer to the soma than in hyperalgesic conditions, and its location was similar to that of the control conditions (Figure 4A). Furthermore, the threshold current of tonically firing SDH neurons recorded from slices 9 - 12 days after CFA injection, when hyperalgesia has resolved, was significantly lower than the threshold current measured in inflammatory conditions and returned to its values measured in the naïve conditions (Figure 4B). In these conditions, all other measured parameters of intrinsic excitability (resting membrane potential, input resistance, AP threshold, AP amplitude and dV/dt_max_) of tonically-firing neurons were similar to those assessed in the naïve or inflammatory conditions (Table 1). These results suggest that the location of the AIS relative to the soma and threshold current of inhibitory SDH neurons change in parallel to the pain sensitivity state.

**Figure 4.**
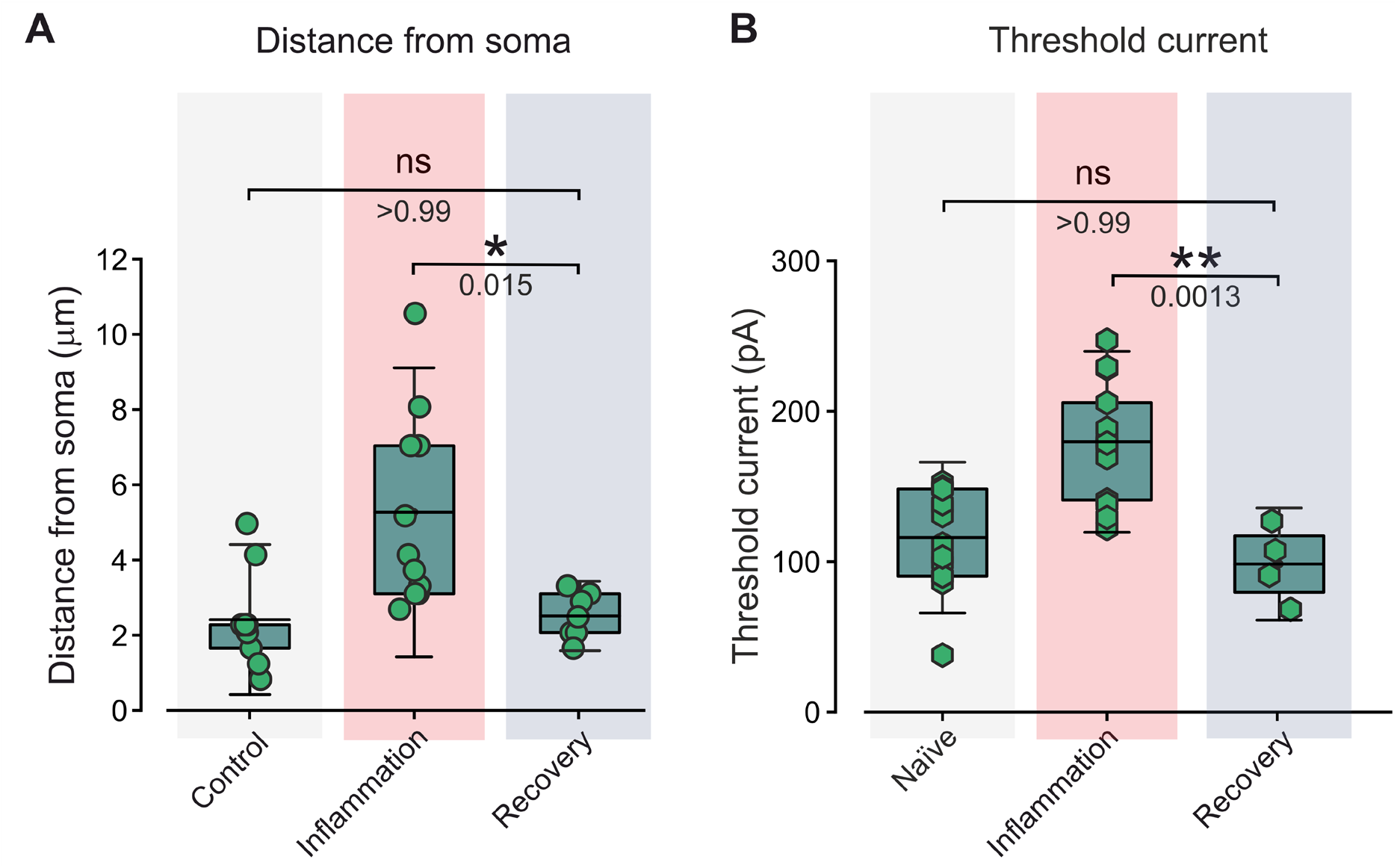
An inflammation-mediated distal shift of AIS and the increase in threshold current reverse when hyperalgesia subsides. **A, B.** Box plots and individual values of the distance of AIS from the soma in Pax2+ neurons (***A***) and threshold current assessed from tonically-firing neurons (***B***) comparing between control conditions (*grey*) and time points at which CFA-mediate hyperalgesia is at its peak (Inflammation, *red*), and CFA-mediated hyperalgesia subsides (Recovery, *purple*). For ***A*** n _Pax2_^+^_, Naive_ = 9 neurons in 9 rats; n _Pax2_^+^_, Inflammation_ = 11 neurons in 11 rats; n _Pax2_^+^_, Recovery_ = 7 neurons in 7 rats, Ordinary one-way ANOVA with posthoc Bonferroni. For ***B*** n _Tonic, Naïve_ = 13 neurons, 13 slices, 7 rats; n _Tonic, Inflammation_ = 14 neurons, 14 slices, 6 rats, n _Tonic, Recovery_ = 4 neurons, 4 slices, 3 rats. Ordinary one-way ANOVA with posthoc Bonferroni. The data for inflammatory and control time points are rearranged from Figures 1E, 1C (AIS distance) and Figure 3B (threshold current).

### A distal shift in the AIS is sufficient to produce an increase in the threshold current

Our results show an inflammation-induced distal shift of the AIS in PAX2^+^ inhibitory neurons concurred with the increase in threshold current in tonically-firing inhibitory neurons, suggesting a link between the AIS shift and change in threshold current. Although previous studies have suggested that an increase in AIS distance from the soma decreases neuronal excitability by increasing the threshold current (Grubb and Burrone, 2010; Rotterman et al., 2021), the opposing relationship between the AIS distance and the level of threshold current was also proposed (Chand et al., 2015; Goethals and Brette, 2020). A possible resolution to these opposing views is the experimental and theoretical data demonstrating that the relation between AIS location and threshold current largely depends on the somatodendritic tree morphology of specific cells (Eyal et al., 2014; Gulledge and Bravo, 2016; Hamada et al., 2016). We, therefore, examined whether, in SDH inhibitory neurons, a distal shift in AIS is sufficient to induce an increase in the threshold current. To this end, we utilized the previously described simplified computational model of a tonically-firing inhibitory neuron (Melnick et al., 2004). We modified this model to emulate the AIS shift from the soma (Figure 5A). Since there are no available data describing the form and conductive properties of the segment between the soma and AIS in SDH inhibitory neurons, we assumed the geometry and conductive properties of this segment as a cone-shaped tapering axon hillock (spacer) which possesses excitable properties similar to the soma (Figure 5A). Implementation of these changes into the model described by Melnick et al. (2004) led to the spontaneous firing of the simulated neuron. We, therefore, adjusted the passive conductance uniformly along the modeled neuron to 5×10^-5^ S cm^−2^. After a change in passive conductance, we recalculated dendritic size and diameter accordingly (*see Methods*). Overall, our model closely resembled the firing behavior of recorded (Figure 5B, Supplementary Figure 1A) or modeled (Melnick et al., 2004) tonically firing neurons. We used this model to measure the change in threshold current as a function of AIS distance from the soma, keeping all other parameters fixed. We found that in these conditions, distancing the AIS from the soma by extending the length of the spacer increased the threshold current (Figure 5C).

**Figure 5.**
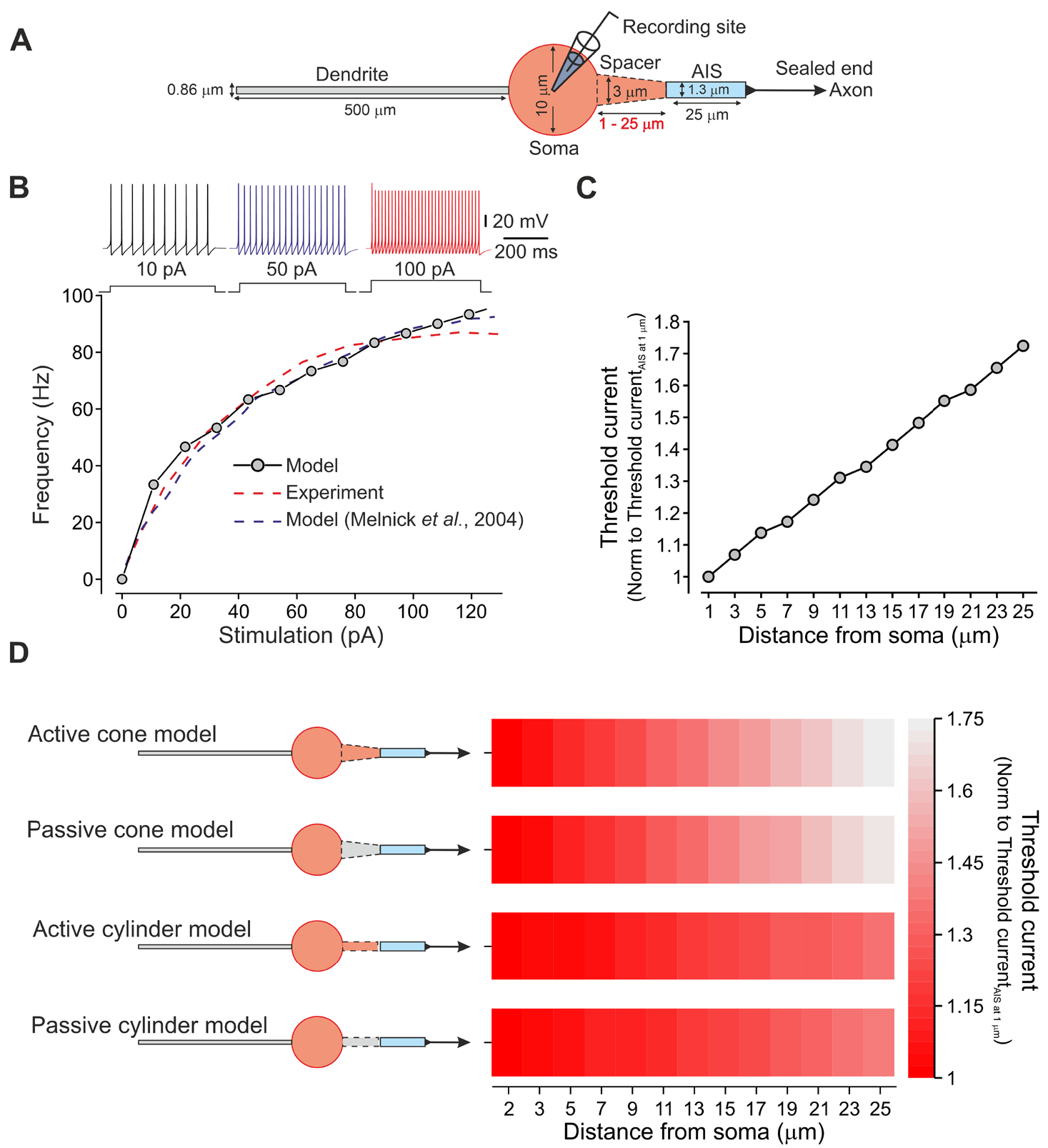
A computational model of SDH tonically-firing neurons predicts that a distal shift in AIS is sufficient to produce an increase in threshold current. **A.** Scheme depicting the general structure of the SDH tonically-firing neuron model. The AIS distance from the soma is varied by the elongation of the cone-shaped spacer. **B.** Modeled frequency-intensity (*f*-I) curves for SDH tonically-firing neuron with an active cone-shaped spacer of 2 μm (*continuous line*) compared with its respective experimental *f*-I curve (*red dashed line*, Supplementary Figure 1B) and *f*-I curve from Melnick et al.’ model of SDH tonically-firing neuron in which the cone-shaped AIS is connected directly to the soma (Melnick et al., 2004, *blue dashed line*). Inset: typical voltage responses to increasing current steps recorded from modeled SDH tonically-firing neuron. **C.** Plot depicting the changes in threshold current with the distance from the soma in the modeled SDH tonically-firing neuron. The threshold current values are normalized to the value measured when AIS is situated 1 μm from the soma (spacer of 1 μm length). **D.** Heat maps of the relation between AIS distance from the soma and the threshold current in the 1 to 25 μm range of distances (*middle*) in various configurations of spacers (shown in *left*): tapered with active conductances (active cone); tapered without active conductances (passive cone); cylindrical with active conductances (active cylinder) and cylindrical without active conductances (passive cylinder). The threshold current values are normalized to the value measured when AIS was situated 1 μm from soma and color-coded (shown on the *right*). Note that AIS distancing leads to an increase in threshold current regardless of the spacer parameters.

To demonstrate that this relation between AIS distance and threshold current does not depend on the geometry and conductive properties of the spacer, we modeled the effect of AIS distance on threshold current for different spacer configurations (Figure 5D). In all examined models (tapering hillock spacer without active conductances and cylinder-shaped spacers with and without active conductance channels), the distancing of AIS away from the soma resulted in an increase in threshold current (Figure 5D). In correlation with previous experimental and theoretical works (Fékété et al., 2021; Goethals and Brette, 2020), demonstrating that higher axonal resistance between soma and AIS increases neuronal excitability, the effect of AIS distancing from the soma on threshold current increase was more pronounced when the spacer was cone-shaped than cylinder-shaped (Figure 5D). The expression of active conductances on a spacer had minimal, if any, effect on the degree of change in threshold current following AIS distancing (Figure 5D).

It is noteworthy that we performed the simulations using the AIS of 25 μm length. There is no data in the literature describing the length of AIS in SDH inhibitory neurons. Our immunohistochemical experiments were not suited for AIS length estimation due to a convoluted axonal trajectory preventing detection of the distal end of the AIS in most cases. We, therefore, chose 25 μm for an AIS length as was proposed for the computational model of SDH inhibitory cell (Melnick et al., 2004). Moreover, 25 μm is the mean of the AIS length variation in computational models, which varies between 10 to 40 μm (Goethals and Brette, 2020) and the described AIS of SC motor neurons and Layer 2,3 and 5 cortical neurons are around 20 - 30 μm (Jamann et al., 2021; Jørgensen et al., 2021; Rotterman et al., 2021). However, to control for a possible effect of AIS length on the AIS distancing-mediated changes on the threshold current, we also performed a series of stimulation with the AIS of 15 μm (Grubb and Burrone, 2010). Simulation of the model by increasing the distancing of the AIS from the soma, using an AIS of 15 μm, also increased the cell threshold current (Supplementary Figure 2). Our computational results suggest that in modeled SDH inhibitory neurons, a distal shift of the AIS over a wide range of distances is sufficient to increase threshold current. These results also imply a link between inflammation-induced AIS distancing and decreased excitability of SDH inhibitory neurons.

## Discussion

Inhibitory and excitatory interneurons located in the SDH receive input from primary nociceptive neurons and process and relay it via spinal cord projection neurons to higher brain centers (Todd, 2010). Thus, the information processing performed by SDH neurons largely determines the output of the SDH networks and establishes pain sensation. In many neuronal systems, the input-output relation of neurons is defined by the molecular and structural properties of the site for the action potential initiation - the AIS. Changes in AIS architecture have been shown to alter neuronal excitability and thereby tune the network output (Grubb et al., 2011; Huang and Rasband, 2018; Jamann et al., 2018). However, the contribution of AIS and its plasticity to the activity of pain-related CNS spinal cord neurons has not yet been determined. Here we provide the first evidence for inflammation-induced AIS plasticity in spinal cord neurons underlying inflammatory hyperalgesia. We show that during inflammatory hyperalgesia, AIS in inhibitory but not excitatory neurons shifts away from the soma, while when hyperalgesia subsides, AIS shifts back close to the soma. This shift in AIS location is accompanied by changes in neuronal excitability, rendering inhibitory neurons less excitable during hyperalgesic inflammation (Figure 6, *right*).

**Figure 6.**
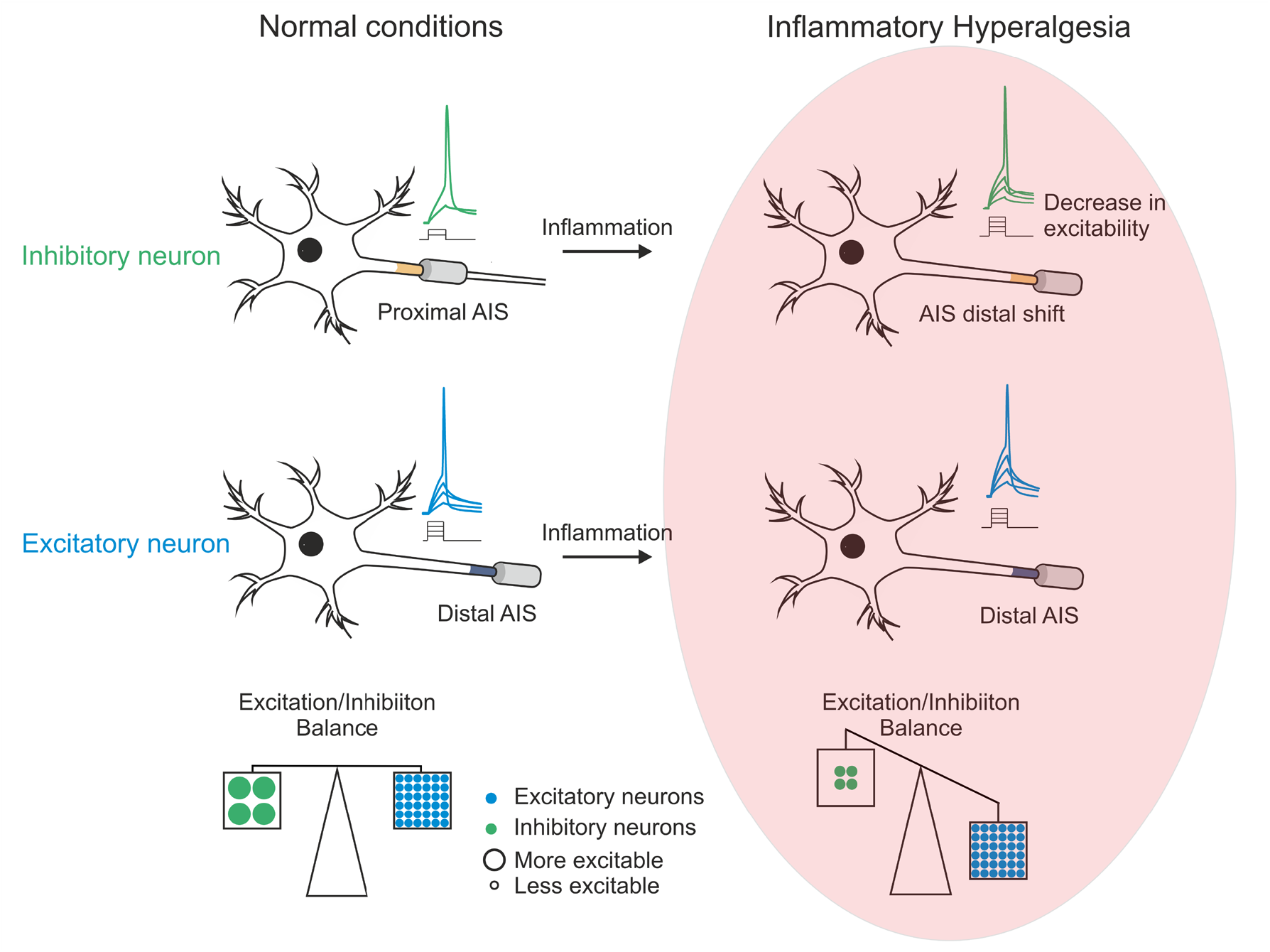
Scheme depicting inflammation-induced AIS plasticity in SDH neurons. *Left*, In normal conditions, the AIS in the inhibitory SDH neurons is located closer to the soma than in excitatory neurons. The proximal location of AIS confers upon SDH inhibitory neurons a small threshold current, suggesting that they will be more prone to activation by the same input than excitatory neurons. This increased intrinsic excitability, together with their amplified “tonic” output, balances the activity of excitatory delayed firing neurons, even though inhibitory neurons only constitute 30% of SDH neurons. *Right*, inflammation induces a distal shift in AIS in inhibitory but not excitatory SDH neurons. The distancing of the AIS from the soma in inhibitory neurons increases the threshold current, which by decreasing their intrinsic excitability, disrupt the excitatory-inhibitory SDH equilibrium, thus rendering the SDH circuitry hyperactive (*blue dots* - excitatory neurons, *green dots* - inhibitory neurons, dot size depicts the level of excitability of the neuron).

We distinguished between inhibitory and excitatory SDH neurons based on the expression of the transcription factor Pax2, promoting inhibitory neuron cell fate (Punnakkal et al., 2014) or neuronal firing pattern. The latter is based on a large body of works (for example, (He et al., 2021; Lee et al., 2019; Punnakkal et al., 2014), showing that the majority of inhibitory neurons fire tonically, while the majority of excitatory interneurons demonstrate delayed firing. The literature that we based our classification on uses a range of current injections showing stable firing patterns (tonic and delayed) together with genetic tools confirming this classification (Tashima et al., 2021). Theoretically, injection of a high enough current could potentially convert delayed firing neurons to tonically firing by overcoming the I_A_ current (Melnick, 2011). However, at the holding potential (−70 mV, see Methods) and at the current steps range we and others have used (Punnakkal et al., 2014), all recorded neurons showed stable firing patterns throughout (Supplementary Figure 1 and Figure 5B, *red dashed line*).

Inhibitory neurons compose a minority (only 30%) of the SDH neurons, and the remaining 70% are excitatory neurons. We show that under naïve conditions, the AIS of inhibitory SDH neurons assumes a position closer to soma than the AIS of excitatory neurons, while the threshold current of inhibitory neurons is lower than that of excitatory neurons, implying that inhibitory neurons are more excitable than the excitatory neurons. This relatively enhanced excitability might compensate for the lower number of inhibitory neurons, balancing the excitation drive from prevalent excitatory neurons (Figure 6, *left*). We also showed that under inflammatory conditions, the AIS of inhibitory neurons assumes a distant location relative to the soma, while the AIS distance from the soma in excitatory neurons did not change its location. Moreover, the threshold current of inhibitory neurons was elevated, while the intrinsic properties of excitatory neurons remained similar to the naïve conditions. This implies that under inflammatory conditions, the excitability of inhibitory neurons is decreased, disrupting the excitatory-inhibitory equilibrium of the SDH circuitry, leading to relative disinhibition (Figure 6, *right*), which, consequently, could increase SDH output towards higher brain centers.

Spinal disinhibition is considered one of the primary mechanisms underlying aberrant sensory processing resulting in pathological pain (Hughes and Todd, 2020). Disruption of the chloride equilibrium or changes in inhibitory synaptic efficacy was proposed as a principal mechanism underlying nerve-injury-mediated pain hypersensitivity (Coull et al., 2005, 2003; Zhang et al., 2018). A decrease in excitability of SDH inhibitory neurons following nerve injury was also demonstrated (Balasubramanyan et al., 2006); however, the underlying biophysical mechanisms have not been identified. Several lines of evidence indicate that spinal disinhibition also contributes to inflammatory pain. COX-2-mediated inhibition of glycine receptors on excitatory neurons was proposed as the mechanism for the inflammation-mediated reduction in spinal inhibition (Zeilhofer and Zeilhofer, 2008). Here we suggest an additional mechanism in which inflammation triggers a distal shift of AIS in inhibitory SDH neurons leading to a decrease in their excitability.

Although the mechanistic link between a distal shift of AIS and a decrease in neuronal excitability has been demonstrated in some neurons (Grubb and Burrone, 2010; Lezmy et al., 2017), this is not a general rule (Chand et al., 2015; Goethals and Brette, 2020). The relation between the AIS distance from the soma and neuronal excitability depends on the relative sizes of the neuronal somatodendritic compartment (Eyal et al., 2014; Gulledge and Bravo, 2016; Hamada et al., 2016). By generating capacitive and conductive loads, the somatodendritic compartment facilitates the escape of inward sodium current from the AIS (Eyal et al., 2014). Thus, in neurons with a large somatodendritic compartment, AIS distancing from the soma leads to electrical isolation of AIS, reducing the conductive and capacitive loads on the AIS, thereby increasing the excitability. On the other hand, AIS distancing from soma increases the dissipation of charges along the axon, thus decreasing excitability. Therefore, the most excitable AIS location along the axon and, therefore, the effect of its shift on neuronal excitability will depend on the balance of AIS’s electrical accessibility to depolarizing drive, generating the threshold current and electrical isolation from somatodendritic conductance loads (Baranauskas et al., 2013; Kuba et al., 2006; Yamada and Kuba, 2016). In order to predict whether the inflammation-mediated AIS distancing we showed in inhibitory neurons could explain an inflammation-induced decrease in their excitability, we used a computational model of inhibitory SDH neurons. It is noteworthy that the model’s purpose was to show the likelihood and the direction of the change in the threshold current following a distal shift in AIS in inhibitory SDH neurons. We did not aim to replicate all the inhibitory neurons’ parameters, considering that the model combines several neuronal morphologies. Thus, our model is not quantitative, but it resembles the tonic output of the inhibitory SDH neurons. Considering these assumptions, our model predicts that in an “averaged” inhibitory SDH neuron, a distal shift of the AIS would be sufficient to cause a decrease in excitability, thus providing a mechanistic link between the distal shift in AIS location and a decrease in the excitability of inhibitory SDH neurons we have observed. The proximal shift in AIS accompanied by the increase in excitability which we have demonstrated after the resolution of inflammatory hyperalgesia, further supports the causal relationship between AIS shift and changes in the excitability of inhibitory SDH neurons. Nevertheless, other factors beyond AIS plasticity (for example, increase in input resistance and increase in functional availability of voltage-gated sodium channels) could affect threshold current. Our data demonstrate that the change in threshold current was not accompanied by changes in other parameters of intrinsic excitability, thus emphasizing the inflammatory-mediated shift in AIS as a key factor in the inflammation-induced decrease in excitability of SDH inhibitory neurons.

By demonstrating the inflammation-mediated shift in AIS location in inhibitory SDH neurons, our results explored only one possible aspect of AIS plasticity affecting neuronal excitability. Differences in channel expression or molecular remodeling of AIS could also affect neuronal excitability (Hinman et al., 2013). In addition, changes in AIS length also modulate neuronal excitability (Gutzmann et al., 2014; Jamann et al., 2021; Kuba et al., 2010). Although several reports show that the changes in AIS length are not correlated with changes in threshold current that we have shown here (Grubb and Burrone, 2010; Rotterman et al., 2021), it is plausible that inflammation affects the molecular composition and length of AIS, thus affecting neuronal excitability via additional mechanisms.

Notably, the inflammation-mediated shift in the AIS we have observed was selective for SDH inhibitory neurons. On a mechanistic level, the shift in AIS location depends on L-type Ca^2+^ channel-induced calcium influx to the AIS (Grubb and Burrone, 2010), triggering calcineurin signaling machinery (Evans et al., 2013). However, single-cell RNA-seq analysis from SDH neurons suggests that L-type calcium channels are expressed mostly by excitatory glutamatergic rather than inhibitory GABAergic SDH neurons (Häring et al., 2018). Following this logic, activation of Cav2.1 (P/Q-type calcium channels) that mediate calcium influx into AIS (Lipkin et al., 2021) and are expressed preferentially in inhibitory SDH neurons (Häring et al., 2018) could be considered as a possible candidate for the inflammation-induced differential AIS shift.

In summary, AIS structural plasticity is a pivotal mechanism underlying changes in neuronal excitability in various pathological conditions such as type 2 diabetes (Yermakov et al., 2018), Alzheimer’s disease (Sun et al., 2014) and stroke (Hinman et al., 2013). Here we show the first evidence of inflammation-induced AIS plasticity in CNS underlying inflammatory hyperalgesia. Moreover, we demonstrate that in pathological pain, the AIS plasticity-mediated network effect might be different from the expected in other neural systems. It has been shown that the increase in input leads to AIS plasticity, resulting in a decrease in neuronal excitability, tuning down the network’s output (Grubb and Burrone, 2010; Jamann et al., 2021; Lezmy et al., 2020, 2017). We show that increased input indeed triggers AIS plasticity and results in a decrease the neuronal excitability, but since it occurs selectively in the inhibitory neurons, the resulting overall network output would be increased. These results are striking in that they demonstrate that inflammation-mediated enhanced input from primary nociceptive neurons can trigger differential changes in the intrinsic excitability of CNS neurons, thus tuning the overall neuronal network response to signal about an ongoing injury. The AIS plasticity, which occurs in a specific neuronal type within the network, provides a novel insight into the effect of this plasticity on network activity.

## Author contributions

Conceptualization, Y.C. and A.M.B.; Methodology Y.C., M.M, Y. K., Y.M. and A.M.B.; Investigation, Y.C., M.M, Y.K., B.K, S.L. and A.M.B.; Writing - Original Draft, Y.C, M.M., B.K, S.L. and A.M.B.; Funding Acquisition, A.M.B.; Supervision, A.M.B.

## Metherials and Methods

### Animals

All procedures were carried out in accordance with the guidelines of the Animal Ethics Committee of the Hebrew University of Jerusalem and were approved by the Ethic committee of The Hebrew University. 5 - 7 week-old Sprague-Dawley (SD) male rats (150 - 225 gr) were housed under controlled temperature (23 ± 2° C) and environment, with ad libitum access to food and water, and kept in a 12-h light/dark cycle.

### Spinal cord slices preparation

Spinal cord slice preparation was performed as previously described by Lu et al. (Lu et al., 2018). In short, rats were anesthetized with isoflurane and decapitated. The vertebral column was quickly removed and immersed in an ice-cold dissecting solution (in mM): 87 NaCl, 2.5 KCl, 1.25 NaH_2_PO_4_, 26 NaHCO_3_, 6 MgCl_2_, 0.5 CaCl_2_, 20 Glucose, 77 sucrose, 1 kynurenic acid, oxygenated with 95/5% O_2_/CO_2_. After nerve roots and meninges were gently removed, the lumbar spinal cord was embedded in low melting agarose (3% in dissecting solution), and parasagittal slices of 300 µm were obtained. Spinal cord slices were then immersed in an oxygenated recovery solution (same as dissecting solution, but without kynurenic acid) at 35°C and allowed to recover for 45 minutes. After 45 minutes, the slices were transferred to an oxygenated storage solution (same as recovery solution, but at room temperature) and until recording.

### Electrophysiology

After recovery, slices were transferred to a submersion recording chamber and mounted on the stage of an upright microscope (BX51WI; Olympus). Slices were then perfused with artificial CSF (aCSF) containing the following (in mM): 125 NaCl, 2.5 KCl, 26 NaHCO_3_, 1.25 NaH_2_PO_4_, and 20 Glucose oxygenated with 95/5% O_2_/CO_2_.

The internal solution was as follows (in mM): 130 K-gluconate, 6 KCl, 8 NaCl, 2 MgATP, and 10 HEPES (pH = 7.4 with KOH). The membrane potential was corrected for a liquid junction potential of +14.9 mV.

Whole-cell current-clamp recordings were performed from neurons located within lamina II, which was identified as a translucent band across the dorsal horn under the microscope. Voltages were recorded in fast current-clamp mode, using a Multiclamp 700B amplifier (Molecular Devices) at room temperature. Data were low-pass filtered at 10 kHz and sampled at 20 kHz for 500 ms current steps protocols and 250 kHz for 10 ms current steps protocols (−3 dB, 8-pole Bessel filter). Patch pipettes were pulled using thick-walled borosilicate glass capillaries (1.5:0.86 mm, outer to the inner diameter) on a P-1000 puller (Sutter Instruments) with a resistance of 3-5 MΩ when filled with standard internal solution (see above), and access resistance was maintained at 7-13 MΩ.

Command current protocols and data digitizing were generated with a Digidata 1440A A/D (Molecular Devices), interfaced with pCLAMP 10.3 (Molecular Devices). Data were analyzed using Clampfit 10.3 software and MATLAB.

For each analyzed SDH neuron first, the firing profile (tonic or delayed) was determined as the patterns of the APs in response to 500-ms depolarizing square pulses (with increments of 0.01 nA). The resting membrane potential (RMP) was maintained at −70 mV in all the analyzed neurons to eliminate the effect of potassium I_A_ current inactivation on the firing pattern (Melnick, 2011). In order to assure the stable firing pattern of the neurons, i.e., that delayed and tonically firing neurons keep their properties regardless of the injected current, all recorded neurons analyzed were classified following measurements at a wide range of current steps.

Apparent input resistance (R_in_) was measured using a series of 500-ms hyperpolarizing and depolarizing square pulses (with increments of 0.01 nA). R_in_ was calculated as the slope of the voltage-current (*V*-I) curve in the linear part of the hyperpolarizing range.

The latency to the first AP was measured following a 500-ms depolarizing pulse of 150 pA, as the time between the beginning of the stimulation step to the peak of the first AP.

Single APs were evoked by 10-ms depolarizing current pulses of 0.01-nA increments. The threshold current (rheobase) was measured as the minimum current required to evoke an action potential (AP). The phase plots of d*V*/d*t* vs. membrane voltage were formed by plotting the rate of change of membrane potential with respect to time (d*V*/d*t*) as a function of membrane potential, following a suprathreshold 10-ms current pulse. Maximum velocities (V/s) were obtained from (d*V*/d*t*)_max_. The AP thresholds were obtained by analyzing the phase plots (d*V*/d*t*) when plotted vs. time and vs. membrane voltage. The threshold voltage was detected using the “first local minimum,” which was determined by analyzing the function from its positive peak to time “0” as the first minimal value of d*V*/d*t* after the peak, followed by an additional increase in the d*V*/d*t*. Then the time of the first local minimum was detected, and its voltage was assessed from the original trace and defined as the threshold voltage.

### Immunohistochemistry

Rats were deeply anesthetized with ketamine and xylazine and fixed by intracardiac perfusion with 20 ml of 4% paraformaldehyde (PFA) freshly prepared in phosphate-buffered saline (PBS, pH 7.4) at 4°C. The lumbar spinal cord was then immediately dissected out, followed by immersion for 1 h in 4% PFA at 4°C, and cryoprotected by overnight immersion in 30% sucrose in PBS at 4°C. Tissue samples were then frozen in OCT medium, and 30 μm cryosections were collected onto Superfrost Plus slides and stored at −20°C.

Slides were first thawed at room temperature (RT), washed in PBS, and incubated for 7 min in a permeabilization solution (0.5% Tween-20 and 1% Triton X-100 in PBS), followed by incubation of 1 h with blocking solution (0.3% Triton X-100, and 5% donkey serum in TBS).

Sections were then incubated overnight at 4°C with primary antibodies, diluted in antibody solution (0.1% Tween-20 and 3% bovine serum albumin in PBS). Then the samples were washed three times for 10 min in PBS, followed by incubation in the dark with fluorescent secondary antibodies, diluted in antibody solution (0.1% Tween-20 and 3% bovine serum albumin in PBS) for 1 h at RT. Finally, sections were washed three times for 10 min in PBS, then dried out and mounted with Vectashield.

#### Antibodies

The triple labeling was performed using (*1*) mouse monoclonal Ankyrin-G (1:250, NeuroMab Facility, Cat# 75-146) with Alexa Fluor 647-conjugated AffiniPure Fab Fragment Goat Anti-Mouse (1:1000, Jackson Immunoresearch, Cat# 115-607-186) as secondary antibody; (*2*) guinea pig polyclonal Anti-NeuN (1:500, Merck, Cat# ABN90P) with Alexa Fluor 488-conjugated AffiniPure Donkey Anti-Guinea Pig (1:1000, Jackson Immunoresearch, Cat# 706-545-148) secondary antibody and (*3*) rabbit polyclonal anti-Pax2 (1:500, Invitrogen, Cat# 71-6000 with DyLight 405-conjugated AffiniPure Goat Anti-Rabbit (1:1000, Jackson Immunoresearch, Cat# 111-475-003) secondary antibody.

#### Image analysis

Images were acquired using a confocal microscope (Axiovert 200 M, Zeiss; Le Pecq, France) and a 60X objective. Images were processed with the NIH ImageJ software (Bethesda, MD, USA). Immunofluorescence intensity profiles were obtained from a segmented line traced along the axon segment on a stack-of-interest projected image using ImageJ. Fluorescence intensity values were normalized to values between zero to one. The distance of the AIS from soma was determined while analyzing Alexa Fluor 647 as the point at which the fluorescence value reached 0.33. Each figure corresponds to a projection image from a *z* stack of 0.5 μm distant optical sections, limited to the object of interest. In order to minimize the measurement errors resulting from the *z* stack processing, we limited the number of maximum stacked sections to 6. Due to this limitation, the length of the AIS itself could not be analyzed because such analysis required much more than 6 sections. In these experiments, the yield of identifying excitatory or inhibitory neurons with labeled AIS was very low, and no more than one neuron per animal was analyzed, such that the numbers indicated in the figures correspond both to the number of neurons and the rats used in the experiments.

### Behavioral experiments

#### Paw withdrawal latency (PWL)

The PWL was measured using Hergeaves apparatus in both hind paws as previously described(Hargreaves et al., 1988). The rats were habituated to the test environment for 1 week in Plexiglas chambers and the experimental surfaces before the behavioral tests. The behavioral baseline was obtained by 3 preliminary measurements. The same investigator performed the scoring in all the behavior tests. Injected solutions were freshly prepared on the day of the experiment.

Animals received a 30-μL intraplantar injection of complete Freund’s adjuvant (CFA; Calbiochem) in the left hind paw. 24 h after CFA injection, the thermal threshold was measured. The behavioral tests were repeated on days 2, 3, 6, 9, 12, and 21.

### Computer simulation

The simplified inhibitory interneuron model was implemented in NEURON 7.8 with the Python interpreter (Dura-Bernal et al., 2019; Hines and Carnevale, 1997). We based our model on a previously published computational model of the SDH inhibitory neuron (Melnick et al., 2004), with adjustments detailed below made to explore the effect of AIS plasticity on cell excitability.

#### Morphology

The morphology of the tonically-firing inhibitory neuron was based on Melnick et al. (Melnick et al., 2004) and included a cylindrical soma (10 µm in diameter, 10 µm in length) connected to an equivalent dendrite segment. The dendrite was 500 µm in length and 0.86 µm in diameter. These parameters were derived from the neuronal properties as described below in “Calculation of dendrite length.” To study the effects of AIS distancing, we modeled an axon hillock which served as a spacer between the soma and the AIS. Since the geometrical and physiological properties of the axon hillock in SDH inhibitory cells were not previously described, we modeled 4 possible axon hillocks configurations: (1) an “active” cone-shaped axon hillock, which decreased gradually from 3 μm in its proximal end to 1.3 μm in the distal end and included somatic active conductances described below; (2) a “passive” cone-shaped axon hillock, which decreased gradually from 3 μm in its proximal end to 1.3 μm in the distal end and included somatic passive conductances described below; (3) an “active” cylindrical axon hillock set to 1.3 μm in diameter and (4) a “passive” cylindrical axon hillock set to 1.3 μm in diameter. The subsequent segment was the AIS compartment with a constant diameter of 1.3 μm which terminated with a sealed-end condition. To simulate changes in AIS location, we gradually increased the length of the axon hillock from 1 to 25 μm while keeping all other aspects of neuron morphology and physiology constant. We used the AIS length of 25 μm as was suggested by (Melnick et al., 2004) and since this length is an average of AIS length variations used in computational models studying AIS (Goethals and Brette, 2020). To rule out the effect of AIS length on the AIS distancing-mediated changes on the threshold current, we also performed a series of stimulation with AIS of 15 μm.

#### Passive membrane properties

Membrane capacitance of 1 μF cm^−2^ was set for all compartments (Melnick et al., 2004). The axial resistance was set at 80 Ω cm (Melnick et al., 2004). In order to prevent spontaneous firing of the cell following changes to axon initial segment morphology, we set the passive membrane resistance to 20000 Ω cm^2^ (passive membrane conductance = 5*10^−5^ S cm^−2^) uniformly for all compartments. The equilibrium potential for passive conductance was −70 mV, and the resting membrane potential was −70 mV (Melnick et al., 2004).

#### Calculation of dendrite length

Following the adjustment of specific passive resistance, the parameters of an equivalent dendrite were calculated using the procedure described previously (Dodge and Cooley, 1973; Rall, 1969, 1959). We used the electronic length of 0.68 and R_in_ of 1.7 GW adapted from (Melnick et al., 2004). We calculated dendrite length and diameter using the parameters described above applied to the equations describing a passive cable:

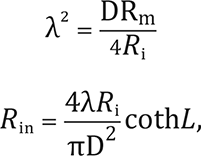

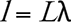

where R_m_ is a membrane resistance, R_i_ is an axial resistance and R_in_ is an input resistance, λ is a characteristic length, *D* is the diameter of the equivalent dendrite, and *l* is the equivalent dendritic length. Accordingly, the calculated dendrite dimensions were: a length of 500 μm and a diameter of 0.86 μm.

#### Active conductances

The model includes sodium (Na) and potassium delayed rectifier (K_DR_) currents as previously described (Melnick et al., 2004) which were fit to Hodgkin Huxley style equations. The steady-state activation variable (*m*_∞_) and the time constant of activation (τ_m_) were determined as *m*_∞_ = α_m_/(α_m_ + β_m_) and τ_m_ = 1/(α_m_ + β_m_) adapted from the original model. For simulating the excitable properties of a neuron, we used the following fixed maximal conductance parameters for each of the segments, maintaining a higher active conductance density at the AIS compared to the remaining neuron.

- gK_DR_ Dendrite = 0.034 S cm^−2^
- gK_DR_ Soma = 0.0043 S cm^−2^
- gK_DR_ AIS = 0.076 S cm^−2^
- gNa Soma = 0.008 S cm^−2^
- gNa AIS = 1.8 S cm^−2^

The reversal potentials for sodium (E_Na_) and potassium (E_K_) were set to +60 mV and −84 mV, respectively. The passive reversal potential was set to −70 mV. All conductances were evenly distributed across the model.

#### Stimulation parameters

All recordings were performed by positioning a NEURON “point-process” electrode at the center of the soma. For computational precision, all compartments were divided into many segments so that the length of individual segments was usually below 1 μm. All simulations were run with 0.05 μs time steps and the nominal temperature of simulations was 23°C.

#### Data analysis

Data generation and plotting were carried out with custom-written Python 2.7 code. To avoid artifacts resulting from the artificial instability of the NEURON simulation environment before reaching steady-state, we allowed for a long 500-ms initialization time before stimulation of the simulated neuron, and this time was discarded from the analysis.

### Statistical analysis

Statistical analyses were performed using Prism 7 (GraphPad). Quantitative data were expressed as the Mean ± SD. We did not perform a power analysis as the proposed experiments are novel, and therefore, we cannot estimate the effect size; however, sample sizes in the current study were similar to previous reports (Lu et al., 2018). Rats were randomly allocated to groups in all experiments. All the analyzed data were normally distributed. The normality was assessed using the Shapiro-Wilk test. Two-tailed unpaired student *t*-tests, ordinary one-way ANOVA and contingency Fisher’s exact test were used when appropriate. Actual *p* values are presented for each data set. The criterion for statistical significance was p < 0.05. Boxplots presented in the figures depict the Mean, 25^th^; 75^th^ percentile, and SD.

### Data and code availability

All datasets generated and/or analyzed during the current study are available in the main text or upon request. The model files are uploaded to Model DB and are also available on request. Further information and requests for resources and reagents should be directed to and will be fulfilled by the Lead Contact, Alexander Binshtok.

## Supplemental Information

**Supplementary Figure 1 (for Figure 2).**
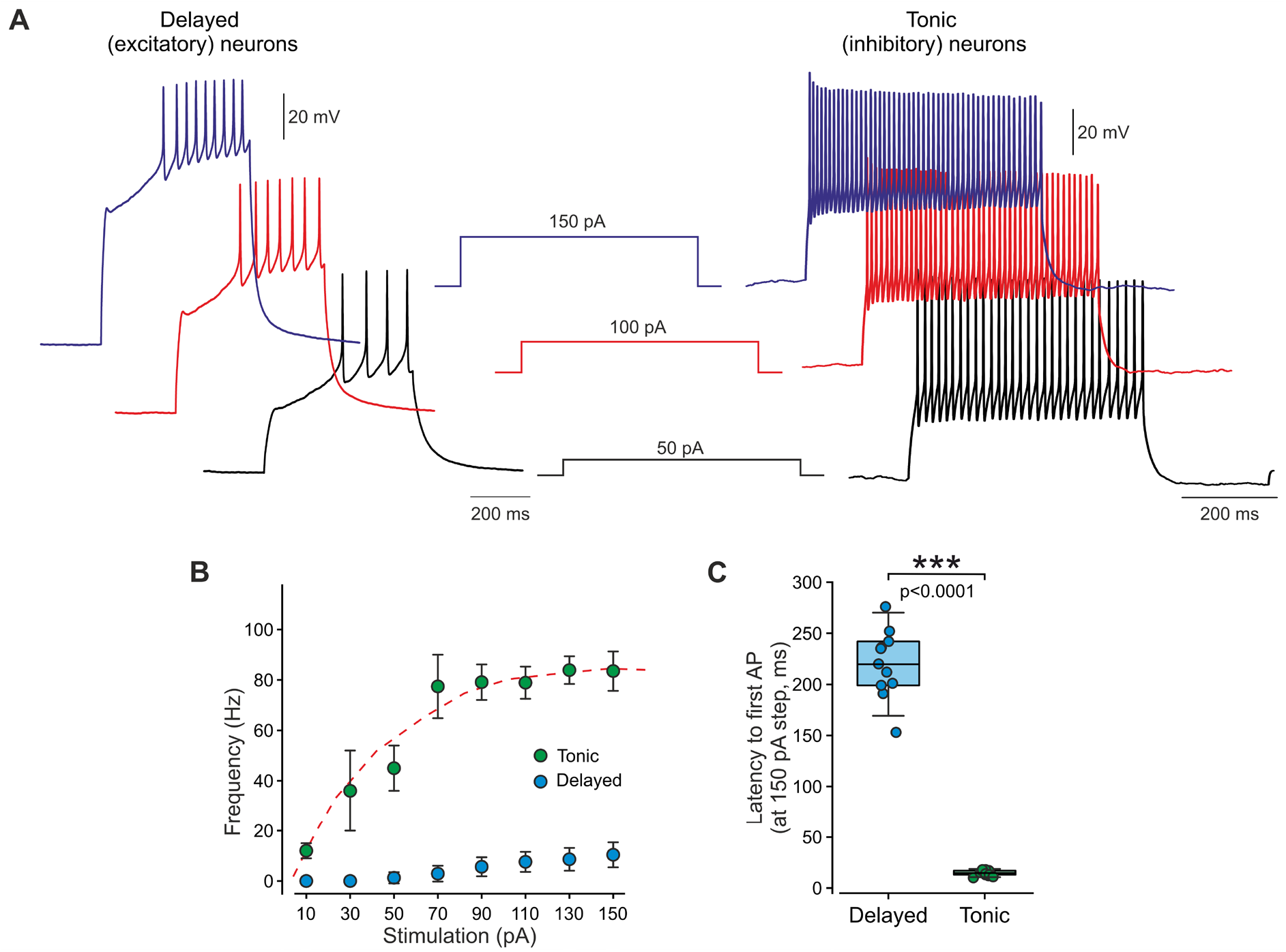
The firing pattern of SDH neurons is stable throughout the increasing stimulation. **A.** An additional example of current-clamp recordings from delayed-firing (“delayed,” *left*) and tonically-firing (“tonic,” *right*) neurons following increasing current steps of 50, 100, and 150 pA demonstrates the stability of the firing properties. Representative of 11 delayed neurons (from 11 slices, 7 rats) and 13 tonic neurons (13 slices, 7 rats). **B.** Mean frequency-intensity (*f*-I) relation and curve (*dashed line*) of tonically-firing (*green*) and delayed-firing (*blue*) neurons. Note that the frequency of delayed-firing neurons remains substantially lower than the tonically-firing neurons even at the highest stimulation steps. Note also the differences in threshold current. n _Delayed_ = 11 neurons, 11 slices, 6 rats; n _Tonic_ = 13 neurons, 13 slices, 7 rats. **C.** Graph comparing box plots and individual values of the latency to the first AP (*see Methods*) in delayed-firing (*blue*) and tonically firing neurons (*green*) following 500-ms step of 150 pA. Note that in all delayed-firing neurons, the latency to the first spike at the maximal stimulating used is significantly delayed compared to the tonically-firing neurons. *** - p < 0.0001; Student’s t-test; n _Delayed_ = 11 neurons, 11 slices, 6 rats; n _Tonic_ = 13 neurons, 13 slices, 7 rats. Box plots depict the mean, 25th, 75th percentile and SD.

**Supplementary Figure 2 (for Figure 5).**
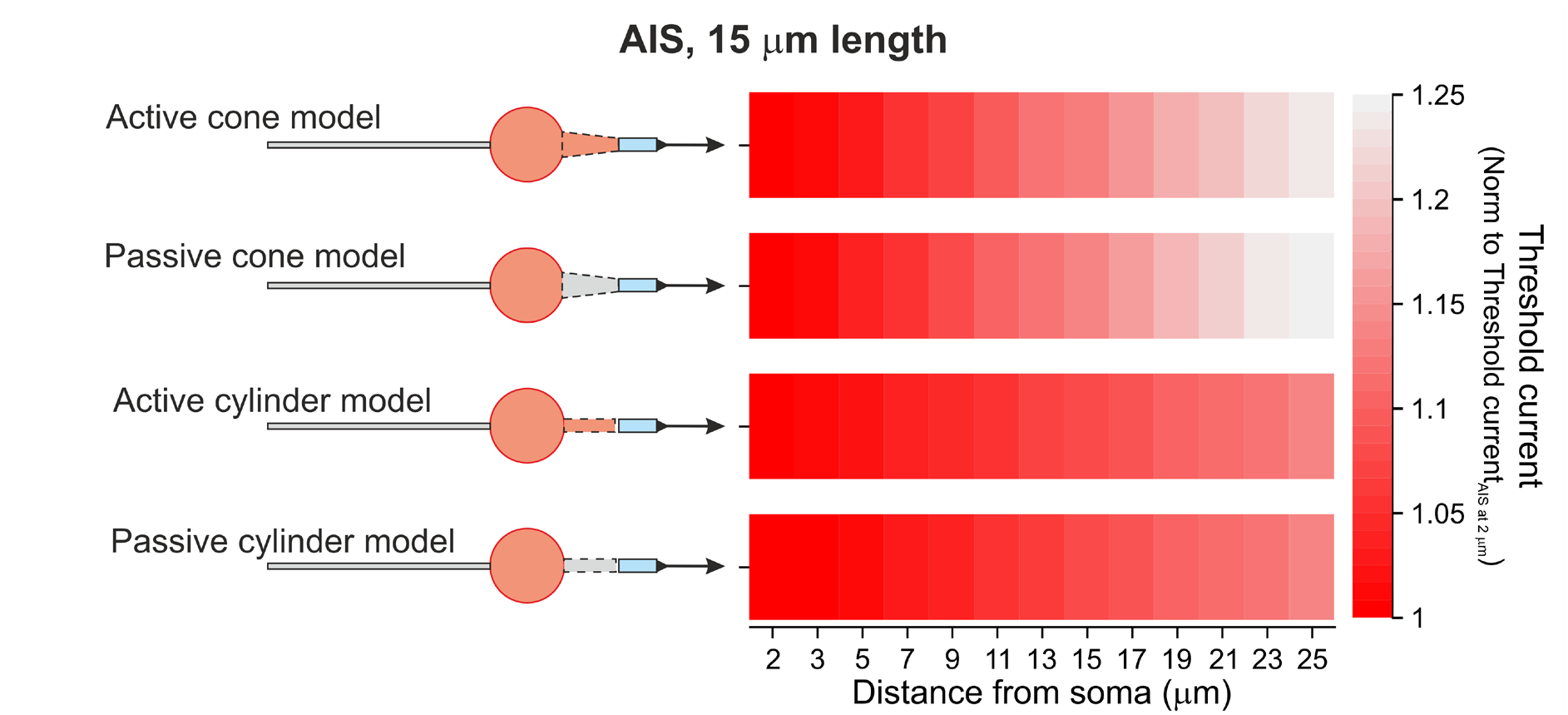
In modeled neurons with a shorter AIS of 15 μm length, the distal shift of AIS leads to an increase in threshold current. Heat maps of the relation between AIS distance from the soma and the threshold current in the 1 to 25 μm range of distances (*middle*) recorded from modeled in SDH tonically-firing neuron with AIS of 15 μm length with various spacers’ configurations (shown in *left*): tapered with active conductances (active cone); tapered without active conductances (passive cone); cylindrical with active conductances (active cylinder) and cylindrical without active conductances (passive cylinder). The threshold current values are normalized to the value measured when AIS was situated 1 μm from soma and color-coded (shown on the *right*). Note that AIS distancing leads to an increase in threshold current regardless of the spacer parameters.

## References

Baba H, Doubell TP, Woolf CJ. 1999. Peripheral Inflammation Facilitates Aβ Fiber-Mediated Synaptic Input to the Substantia Gelatinosa of the Adult Rat Spinal Cord. Journal of Neuroscience 19:859–867. doi:10.1523/JNEUROSCI.19-02-00859.1999

Balasubramanyan S, Stemkowski PL, Stebbing MJ, Smith PA. 2006. Sciatic chronic constriction injury produces cell-type-specific changes in the electrophysiological properties of rat substantia gelatinosa neurons. Journal of Neurophysiology 96:579–590. doi:10.1152/jn.00087.2006

Baranauskas G, David Y, Fleidervish IA. 2013. Spatial mismatch between the Na+ flux and spike initiation in axon initial segment. Proc Natl Acad Sci U S A 110:4051–4056. doi:10.1073/pnas.1215125110

Barkai O, Butterman R, Katz B, Lev S, Binshtok AM. 2020. The Input-Output Relation of Primary Nociceptive Neurons is Determined by the Morphology of the Peripheral Nociceptive Terminals. J Neurosci 40:9346–9363. doi:10.1523/JNEUROSCI.1546-20.2020

Barkai O, Puig S, Lev S, Title B, Katz B, Eli-Berchoer L, Gutstein HB, Binshtok AM. 2019. Platelet-derived growth factor activates nociceptive neurons by inhibiting M-current and contributes to inflammatory pain. Pain 160:1281–1296. doi:10.1097/j.pain.0000000000001523

Basbaum AI, Bautista DM, Scherrer GG, Julius D. 2009. Cellular and Molecular Mechanisms of Pain. Cell 139:267–284. doi:10.1016/j.cell.2009.09.028

Battefeld A, Tran BT, Gavrilis J, Cooper EC, Kole MH. 2014. Heteromeric Kv7.2/7.3 channels differentially regulate action potential initiation and conduction in neocortical myelinated axons. J Neurosci 34:3719–3732. doi:10.1523/JNEUROSCI.4206-13.2014

Bean BP. 2007. The action potential in mammalian central neurons. Nat Rev Neurosci 8. doi:10.1038/nrn2148

Bender KJ, Trussell LO. 2012. The physiology of the axon initial segment. Annu Rev Neurosci 35:249–265. doi:10.1146/ANNUREV-NEURO-062111-150339

Binshtok AM, Wang H, Zimmermann K, Amaya F, Vardeh D, Shi L, Brenner GJ, Ji R-RR, Bean BP, Woolf CJ, Samad TA. 2008. Nociceptors are interleukin-1beta sensors. J Neurosci 28:14062–14073.

Braz J, Solorzano C, Wang X, Basbaum AI. 2014. Transmitting Pain and Itch Messages: A Contemporary View of the Spinal Cord Circuits that Generate Gate Control. Neuron. doi:10.1016/j.neuron.2014.01.018

Buffington SA, Rasband MN. 2011. The axon initial segment in nervous system disease and injury. European Journal of Neuroscience 34:1609–1619. doi:10.1111/j.1460-9568.2011.07875.x

Chand AN, Galliano E, Chesters RA, Grubb MS. 2015. A Distinct Subtype of Dopaminergic Interneuron Displays Inverted Structural Plasticity at the Axon Initial Segment. Journal of Neuroscience 35:1573–1590. doi:10.1523/JNEUROSCI.3515-14.2015

Coull JAM, Beggs S, Boudreau D, Boivin D, Tsuda M, Inoue K, Gravel C, Salter MW, de Koninck Y. 2005. BDNF from microglia causes the shift in neuronal anion gradient underlying neuropathic pain. Nature 438:1017–1021. doi:10.1038/nature04223

Coull JAM, Boudreau D, Bachand K, Prescott SA, Nault F, Sík A, de Koninck P, de Koninck Y. 2003. Trans-synaptic shift in anion gradient in spinal lamina I neurons as a mechanism of neuropathic pain. Nature 424:938–942. doi:10.1038/nature01868

Devor M. 2006. Sodium channels and mechanisms of neuropathic pain. The journal of pain : official journal of the American Pain Society 7:S3–S12.

Djouhri L, Koutsikou S, Fang X, McMullan S, Lawson SN. 2006. Spontaneous pain, both neuropathic and inflammatory, is related to frequency of spontaneous firing in intact C-fiber nociceptors. J Neurosci 26:1281–1292. doi:10.1523/jneurosci.3388-05.2006

Dodge FA, Cooley JW. 1973. Action potential of the motorneuron. IBM Journal of Research and Development 17:219–229. doi:10.1147/RD.173.0219

Dura-Bernal S, Suter BA, Gleeson P, Cantarelli M, Quintana A, Rodriguez F, Kedziora DJ, Chadderdon GL, Kerr CC, Neymotin SA, McDougal RA, Hines M, Shepherd GM, Lytton WW. 2019. NetPyNE, a tool for data-driven multiscale modeling of brain circuits. Elife 8. doi:10.7554/ELIFE.44494

Evans MD, Sammons RP, Lebron S, Dumitrescu AS, Watkins TBK, Uebele VN, Renger JJ, Grubb MS. 2013. Calcineurin Signaling Mediates Activity-Dependent Relocation of the Axon Initial Segment. Journal of Neuroscience 33:6950–6963. doi:10.1523/JNEUROSCI.0277-13.2013

Eyal G, Mansvelder HD, de Kock CPJ, Segev I. 2014. Dendrites Impact the Encoding Capabilities of the Axon. Journal of Neuroscience 34:8063–8071. doi:10.1523/JNEUROSCI.5431-13.2014

Fékété A, Ankri N, Brette R, Debanne D. 2021. Neural excitability increases with axonal resistance between soma and axon initial segment. Proc Natl Acad Sci U S A 118. doi:10.1073/PNAS.2102217118/SUPPL_FILE/PNAS.2102217118.SAPP.PDF

Galliano E, Hahn C, Browne LP, Villamayor PR, Tufo C, Crespo A, Grubb MS. 2021. Brief Sensory Deprivation Triggers Cell Type-Specific Structural and Functional Plasticity in Olfactory Bulb Neurons. Journal of Neuroscience 41:2135–2151. doi:10.1523/JNEUROSCI.1606-20.2020

Goethals S, Brette R. 2020. Theoretical relation between axon initial segment geometry and excitability. Elife 9. doi:10.7554/eLife.53432

Goldstein RH, Barkai O, Íñigo-Portugués A, Katz B, Lev S, Binshtok AM. 2019. Location and Plasticity of the Sodium Spike Initiation Zone in Nociceptive Terminals In Vivo. Neuron 102:801–812.

Grubb MS, Burrone J. 2010. Activity-dependent relocation of the axon initial segment fine-tunes neuronal excitability. Nature 465:1070–1074. doi:10.1038/nature09160

Grubb MS, Shu Y, Kuba H, Rasband MN, Wimmer VC, Bender KJ. 2011. Short-and long-term plasticity at the axon initial segment. Journal of Neuroscience 31:16049–16055. doi:10.1523/JNEUROSCI.4064-11.2011

Gudes S, Barkai O, Caspi Y, Katz B, Lev S, Binshtok AM. 2015. The role of slow and persistent TTX-resistant sodium currents in acute tumor necrosis factor-alpha-mediated increase in nociceptors excitability. J Neurophysiol 113:601–619. doi:10.1152/jn.00652.2014

Gulledge AT, Bravo JJ. 2016. Neuron Morphology Influences Axon Initial Segment Plasticity. eNeuro 3:255–265. doi:10.1523/ENEURO.0085-15.2016

Gutzmann A, Ergül N, Grossmann R, Schultz C, Wahle P, Engelhardt M. 2014. A period of structural plasticity at the axon initial segment in developing visual cortex. Frontiers in Neuroanatomy 8:11. doi:10.3389/FNANA.2014.00011/ABSTRACT

Ha SR, Rasband MN. 2019. The SIZ of Pain. Neuron. doi:10.1016/j.neuron.2019.04.040

Hamada MS, Goethals S, de Vries SI, Brette R, Kole MHP. 2016. Covariation of axon initial segment location and dendritic tree normalizes the somatic action potential. Proc Natl Acad Sci U S A 113:14841–14846. doi:10.1073/PNAS.1607548113/-/DCSUPPLEMENTAL

Hargreaves K, Dubner R, Brown F, Flores C, Joris J. 1988. A new and sensitive method for measuring thermal nociception in cutaneous hyperalgesia. Pain 32:77–88.

Häring M, Zeisel A, Hochgerner H, Rinwa P, Jakobsson JET, Lönnerberg P, la Manno G, Sharma N, Borgius L, Kiehn O, Lagerström MC, Linnarsson S, Ernfors P. 2018. Neuronal atlas of the dorsal horn defines its architecture and links sensory input to transcriptional cell types. Nature Neuroscience 2018 21:6 21:869–880. doi:10.1038/s41593-018-0141-1

He X, Liu P, Zhang X, Jiang Z, Gu N, Wang Q, Lu Y. 2021. Molecular and Electrophysiological Characterization of Dorsal Horn Neurons in a GlyT2-iCre-tdTomato Mouse Line. J Pain Res 14:907–921. doi:10.2147/JPR.S296940

Hines ML, Carnevale NT. 1997. The NEURON simulation environment. Neural Comput 9:1179–1209.

Hinman JD, Rasband MN, Carmichael ST. 2013. Remodeling of the axon initial segment after focal cortical and white matter stroke. Stroke 44:182–189. doi:10.1161/STROKEAHA.112.668749

Huang CY-M, Rasband MN. 2018. Axon initial segments: structure, function, and disease. Ann N Y Acad Sci 1420:46–61. doi:10.1111/NYAS.13718

Hucho T, Levine JD. 2007. Signaling pathways in sensitization: toward a nociceptor cell biology. Neuron 55:365–376.

Hughes DI, Todd AJ. 2020. Central Nervous System Targets: Inhibitory Interneurons in the Spinal Cord. Neurotherapeutics. doi:10.1007/s13311-020-00936-0

Jamann N, Dannehl D, Lehmann N, Wagener R, Thielemann C, Schultz C, Staiger J, Kole MHP, Engelhardt M. 2021. Sensory input drives rapid homeostatic scaling of the axon initial segment in mouse barrel cortex. Nature Communications 12. doi:10.1038/S41467-020-20232-X

Jamann N, Jordan M, Engelhardt M. 2018. Activity-Dependent Axonal Plasticity in Sensory Systems. Neuroscience 368:268–282. doi:10.1016/J.NEUROSCIENCE.2017.07.035

Jenerick H. 1963. Phase plane trajectories of the muscle spike potential. Biophys J 3:363–377.

Jin X, Gereau RW th. 2006. Acute p38-mediated modulation of tetrodotoxin-resistant sodium channels in mouse sensory neurons by tumor necrosis factor-alpha. J Neurosci 26:246–255.

Jørgensen HS, Jensen DB, Dimintiyanova KP, Bonnevie VS, Hedegaard A, Lehnhoff J, Moldovan M, Grondahl L, Meehan CF. 2021. Increased Axon Initial Segment Length Results in Increased Na+ Currents in Spinal Motoneurones at Symptom Onset in the G127X SOD1 Mouse Model of Amyotrophic Lateral Sclerosis. Neuroscience 468:247–264. doi:10.1016/J.NEUROSCIENCE.2020.11.016

Kohno T, Moore KA, Baba H, Woolf CJ. 2003. Peripheral nerve injury alters excitatory synaptic transmission in lamina II of the rat dorsal horn. J Physiol 548:131–138.

Kole MH, Brette R. 2018. The electrical significance of axon location diversity. Current Opinion in Neurobiology. doi:10.1016/j.conb.2018.02.016

Kole MHP, Ilschner SU, Kampa BM, Williams SR, Ruben PC, Stuart GJ. 2008. Action potential generation requires a high sodium channel density in the axon initial segment. Nature Neuroscience 11:178–186. doi:10.1038/NN2040

Kole MHP, Stuart GJ. 2012. Signal Processing in the Axon Initial Segment. Neuron 73:235–247. doi:10.1016/J.NEURON.2012.01.007

Kordeli E, Lambert S, Bennett V. 1995. AnkyrinG. A new ankyrin gene with neural-specific isoforms localized at the axonal initial segment and node of Ranvier. J Biol Chem 270:2352–2359. doi:10.1074/JBC.270.5.2352

Kuba H, Ishii TM, Ohmori H. 2006. Axonal site of spike initiation enhances auditory coincidence detection. Nature 444:1069–1072. doi:10.1038/nature05347

Kuba H, Oichi Y, Ohmori H. 2010. Presynaptic activity regulates Na(+) channel distribution at the axon initial segment. Nature 465:1075–1078.

Lee KY, Ratté S, Prescott SA. 2019. Excitatory neurons are more disinhibited than inhibitory neurons by chloride dysregulation in the spinal dorsal horn. Elife 8. doi:10.7554/ELIFE.49753

Lennertz RC, Kossyreva EA, Smith AK, Stucky CL. 2012. TRPA1 mediates mechanical sensitization in nociceptors during inflammation. PLoS One 7:23. doi:10.1371/journal.pone.0043597

Leterrier C. 2018. The Axon Initial Segment: An Updated Viewpoint. Journal of Neuroscience 38:2135–2145. doi:10.1523/JNEUROSCI.1922-17.2018

Lezmy J, Gelman H, Katsenelson M, Styr B, Tikochinsky E, Lipinsky M, Peretz A, Slutsky I, Attali B. 2020. M-Current Inhibition in Hippocampal Excitatory Neurons Triggers Intrinsic and Synaptic Homeostatic Responses at Different Temporal Scales. Journal of Neuroscience 40:3694–3706. doi:10.1523/JNEUROSCI.1914-19.2020

Lezmy J, Lipinsky M, Khrapunsky Y, Patrich E, Shalom L, Peretz A, Fleidervish IA, Attali B. 2017. M-current inhibition rapidly induces a unique CK2-dependent plasticity of the axon initial segment. Proc Natl Acad Sci U S A 114:E10234–e10243. doi:10.1073/pnas.1708700114

Li J, Kritzer E, Craig PE, Baccei ML. 2015. Aberrant Synaptic Integration in Adult Lamina I Projection Neurons Following Neonatal Tissue Damage. Journal of Neuroscience. doi:10.1523/JNEUROSCI.3585-14.2015

Lipkin AM, Cunniff MM, Spratt PWE, Lemke SM, Bender KJ. 2021. Functional Microstructure of CaV-Mediated Calcium Signaling in the Axon Initial Segment. Journal of Neuroscience 41:3764–3776. doi:10.1523/JNEUROSCI.2843-20.2021

Lorincz A, Nusser Z. 2008. Cell-Type-Dependent Molecular Composition of the Axon Initial Segment. Journal of Neuroscience 28:14329–14340. doi:10.1523/JNEUROSCI.4833-08.2008

Lu Y, Doroshenko M, Lauzadis J, Kanjiya MP, Rebecchi MJ, Kaczocha M, Puopolo M. 2018. Presynaptic Inhibition of Primary Nociceptive Signals to Dorsal Horn Lamina I Neurons by Dopamine. Journal of Neuroscience 38:8809–8821. doi:10.1523/JNEUROSCI.0323-18.2018

Melnick I v. 2011. Voltage-Dependent Behavior of Delayed-Firing Neurons in the Rat Substantia Gelatinosa. Neurophysiology 42:287–293.

Melnick I v., Santos SFA, Szokol K, Szûcs P, Safronov B v. 2004. Ionic Basis of Tonic Firing in Spinal Substantia Gelatinosa Neurons of Rat. Journal of Neurophysiology 91:646–655. doi:10.1152/jn.00883.2003

Punnakkal P, Schoultz C von, Haenraets K, Wildner H, Zeilhofer HU. 2014. Morphological, biophysical and synaptic properties of glutamatergic neurons of the mouse spinal dorsal horn. The Journal of Physiology 592:759–776. doi:10.1113/JPHYSIOL.2013.264937

Rall W. 1969. Time Constants and Electrotonic Length of Membrane Cylinders and Neurons. Biophysical Journal 9:1483–1508. doi:10.1016/S0006-3495(69)86467-2

Rall W. 1959. Branching dendritic trees and motoneuron membrane resistivity. Exp Neurol 1:491–527. doi:10.1016/0014-4886(59)90046-9

Rotterman TM, Carrasco DI, Housley SN, Nardelli P, Powers RK, Cope TC. 2021. Axon initial segment geometry in relation to motoneuron excitability. PLoS One 16. doi:10.1371/JOURNAL.PONE.0259918

Sasaki T. 2013. The axon as a unique computational unit in neurons. Neuroscience Research 75:83–88. doi:10.1016/J.NEURES.2012.12.004

Shevchuk DP, Agashkov KS, Bilan P v., Voitenko N v. 2017. Spontaneous Synaptic Activity in Projection Neurons of Lamina I of the Isolated Rat Lumbar Spinal Cord: Effect of Peripheral Inflammation. Neurophysiology 2017 49:4 49:301–304. doi:10.1007/S11062-017-9686-Y

Sorkin LS, Xiao WH, Wagner R, Myers RR. 1997. Tumour necrosis factor-alpha induces ectopic activity in nociceptive primary. Neuroscience 81:255–262.

Sun X, Wu Y, Gu M, Liu Z, Ma Y, Li J, Zhang Y. 2014. Selective filtering defect at the axon initial segment in Alzheimer’s disease mouse models. Proc Natl Acad Sci U S A 111:14271–14276. doi:10.1073/PNAS.1411837111/-/DCSUPPLEMENTAL

Tal M, Devor M. 1992. Ectopic discharge in injured nerves: comparison of trigeminal and somatic afferents. Brain Res 579:148–151.

Tashima R, Koga K, Yoshikawa Y, Sekine M, Watanabe M, Tozaki-Saitoh H, Furue H, Yasaka T, Tsuda M. 2021. A subset of spinal dorsal horn interneurons crucial for gating touch-evoked pain-like behavior. Proc Natl Acad Sci U S A 118. doi:10.1073/PNAS.2021220118/SUPPL_FILE/PNAS.2021220118.SAPP.PDF

Telenczuk M, Fontaine B, Brette R. 2017. The basis of sharp spike onset in standard biophysical models. PLOS ONE 12:e0175362. doi:10.1371/JOURNAL.PONE.0175362

Todd AJ. 2010. Neuronal circuitry for pain processing in the dorsal horn. Nature Reviews Neuroscience 11:823–836. doi:10.1038/nrn2947

Wercberger R, Basbaum AI. 2019. Spinal cord projection neurons: a superficial, and also deep analysis. Current Opinion in Physiology 11:109–115. doi:10.1016/j.cophys.2019.10.002

Yamada R, Kuba H. 2016. Structural and Functional Plasticity at the Axon Initial Segment. Front Cell Neurosci 10.

Yarmolinsky DA, Peng Y, Pogorzala LA, Rutlin M, Hoon MA, Zuker CS. 2016. Coding and Plasticity in the Mammalian Thermosensory System. Neuron 92:1079–1092. doi:10.1016/j.neuron.2016.10.021

Yermakov LM, Drouet DE, Griggs RB, Elased KM, Susuki K. 2018. Type 2 diabetes leads to axon initial segment shortening in db/db mice. Frontiers in Cellular Neuroscience 12:146. doi:10.3389/FNCEL.2018.00146/BIBTEX

Zeilhofer HU, Zeilhofer UB. 2008. Spinal dis-inhibition in inflammatory pain. Neuroscience Letters 437:170–174. doi:10.1016/j.neulet.2008.03.056

Zhang H, Li Y, Yang Q, Liu XG, Dougherty PM. 2018. Morphological and physiological plasticity of spinal lamina II GABA neurons is induced by sciatic nerve chronic constriction injury in mice. Frontiers in Cellular Neuroscience 12. doi:10.3389/fncel.2018.00143

